# Zebrafish caudal fin amputation induces a metabolic switch necessary for cell identity transitions and cell cycle re-entry to support blastema formation and bone regeneration

**DOI:** 10.1101/2022.01.26.477895

**Authors:** Ana S. Brandão, Jorge Borbinha, Telmo Pereira, Patrícia H. Brito, Raquel Lourenço, Anabela Bensimon-Brito, António Jacinto

**Affiliations:** CEDOC, NOVA Medical School, NOVA University of Lisbon, Campo Mártires da Pátria 130, Lisboa 1169-056, Portugal; UCIBIO, Dept. Ciências da Vida, Faculdade de Ciências e Tecnologia, Universidade NOVA de Lisboa, 2819-516 Caparica, Portugal; i4HB, Associate Laboratory - Institute for Health and Bioeconomy, Faculdade de Ciências e Tecnologia, Universidade NOVA de Lisboa, 2819-516 Caparica, Portugal; INSERM, ATIP-Avenir, Aix Marseille Univ, Marseille Medical Genetics, Marseille, France

## Abstract

Regeneration depends on the ability of mature cells at the injury site to respond to injury, generating tissue-specific progenitors that incorporate the blastema and proliferate to reconstitute the original organ architecture. The metabolic microenvironment has been tightly connected to cell function and identity during development and tumorigenesis. Yet, the link between metabolism and cell identity at the mechanistic level in a regenerative context remains unclear. The adult zebrafish caudal fin, and bone cells specifically, have been crucial for the understanding of mature cell contribution to tissue regeneration. Here, we use this model to explore the relevance of glucose metabolism for the cell fate transitions preceding new osteoblast formation and blastema assembly. We show that injury triggers a shift in the metabolic profile at early stages of regeneration, enhancing glycolysis at the expense of mitochondrial oxidation. This metabolic switch mediates transcriptional changes that make mature osteoblast amenable to be reprogramed into pre-osteoblasts and induces cell cycle re-entry and progression. Manipulation of the metabolic profile led to severe reduction of the pre-osteoblast pool, diminishing their capacity to generate new osteoblasts, and to a complete abrogation of blastema formation. Overall, our data indicate that metabolic alterations have a powerful instructive role in regulating genetic programs that dictate fate decisions and stimulate proliferation, thereby providing a deeper understanding on the mechanisms regulating blastema formation and bone regeneration.

## Introduction

The zebrafish *(Danio rerio)* is a well-established model to study vertebrate regeneration due to its capacity to efficiently regenerate multiple organs and complex tissues, such as the caudal fin(1–3). Caudal fin regeneration is an epimorphic process dependent on the formation of a blastema, a transient structure composed of proliferative lineage-restricted progenitor cells that originate from mature cells of the uninjured tissue(4, 5). One of the main components of the caudal fin are the segmented bone elements, or bony-rays(6, 7), which have been shown to regenerate from new osteoblasts (OBs) arising from dedifferentiation of mature OBs close to the amputation site. Mature OBs acquire the cellular properties of less differentiated cells or pre-osteoblasts (pre-OBs)(8–11), which proliferate and redifferentiate into mature bone-producing OBs, thereby reconstituting the fin skeletal tissue(12–15). Mammals, in contrast to zebrafish, have a poor ability to regenerate(16–18) . Mammalian bone, in particular, has an intrinsic capacity to be remodelled throughout life and, to a certain extent, repair after fracture(19, 20), but it fails to fully regenerate. This is mainly achieved through activation of signalling cascades that culminate in the recruitment and differentiation of osteoprogenitors into OBs, the bone-forming cells. In this scenario, understanding how OBs and OB sources are activated and recruited in zebrafish can greatly contribute to new therapeutic solutions to improve bone formation and restoration in mammals(21). Activation of reprogramming events that allow cells to change fate in response to injury appear to be conserved biological processes, which are used to prompt tissue regeneration in several organisms(18,22,23), including in mammals(24, 25). Therefore, it is crucial to identify the early molecular events inducing cell dedifferentiation to promote tissue regeneration.

Cell identity and the functional state often reflect a specific metabolic profile that depends, for instance, on nutrient and oxygen availability, and on bioenergetics and biomass requirements(26, 27). Metabolic routes serve not only the crucial purpose of converting or use energy to maintain cellular integrity and survival, but also have a pivotal role in restructuring gene expression to determine cell identity and function by influencing cell signalling and epigenetic modulators(28, 29). Glucose metabolism is currently regarded as a powerful instructor of cell fate decisions(30–32) during development. It is well documented that stem cells and their differentiated progeny have distinct metabolic preferences: stem cells often favour non-oxidative glycolysis, while differentiated non-dividing somatic cells rely on mitochondrial oxidative phosphorylation (OXPHOS) (33–36). Importantly, the metabolic signature is not static and can rapidly switch according to the cellular demands, a phenomenon commonly designated as metabolic reprogramming(26, 32). This is particularly relevant under certain environmental conditions when tissue homeostasis is breached, such as during inflammation, and during disease. Both activated T cells and pro-inflammatory macrophages switch metabolic profiles compatible with a prevalence of aerobic glycolysis(37, 38). The shift to aerobic glycolysis is also observed in cancer cells, where it is referred as the Warburg effect(39–43).

Despite the considerable amount of information on the role of metabolism during development and disease(43–47), the link between glucose metabolism and regeneration is still far from understood. Shifts in metabolism leading to a prevalence of glycolysis have already been observed in planarians and amphibians during regeneration (48–51). More recently, glycolysis was proposed to modulate specific aspects of regeneration in the zebrafish, specifically in the larval tail(52) and in the heart(53, 54). Nevertheless, much remains to be elucidated on how metabolism influences changes in cell identity necessary to promote tissue regeneration.

Here, we use the adult zebrafish caudal fin to determine the role of energy metabolism in mature OB dedifferentiation and pre-OB recruitment. By performing a series of gene expression and metabolomic studies, we observed that changes in the metabolic signature triggered upon amputation are at the core of the initial set-up of the cellular programmes that control caudal fin regeneration. These data indicate that mature OBs, and possibly other cell lineages in the caudal fin, undergo a metabolic reprogramming, prioritizing glycolysis in detriment of OXPHOS. We further demonstrate that this switch is necessary for mature OBs to commit to a specialized genetic program that enables them to dedifferentiate, proliferate and act as progenitor cells, ensuring proper bone regeneration. Taken together, our results demonstrate that metabolic reprogramming is one of the earliest described cellular events that dictate adult caudal fin and bone regeneration.

## Results

### 1. Osteoblast dedifferentiation occurs before blastema formation and during the wound healing phase

Caudal fin bone regeneration is achieved through activation of cell sources, such as mature OBs, that change their identity to generate new OBs. Under homeostatic conditions, mature OBs reside as quiescent bone-lining cells, but lose their differentiated character when undergoing dedifferentiation(8, 9). This process is characterized by the downregulation of mature markers, such as *bone gamma-carboxyglutamic acid-containing protein* (*bglap*) at 12 hpa (8,9,55), and upregulation of the pre-OB marker, *runx2,* at 24 hpa (8,9,19). OBs undergoing dedifferentiation detach from the bony-ray surface via an EMT-like event, migrate towards the stump region, re-enter the cell cycle and form a pool of pre-OB cells that incorporates the blastema (Fig 1A)(8,9,12). To determine if OBs show signs of dedifferentiation in early stages of regeneration, we increase the time resolution of the initial OB response to amputation. First, we analyse the relative expression of mature (*bglap*) and pre-OB (*runx2a, runx2b*) markers 6 hours post-amputation (hpa) in regenerating caudal fins in relation to uninjured caudal fins (0 hpa). We observed a downregulation of *bglap* and upregulation of *runx2a* (Fig 1B) suggesting that, in contrast to previous studies(8, 9), mature OBs are already undergoing transcriptional changes consistent with dedifferentiation as early as 6 hpa. We then analysed the *bglap*:EGFP transgenic line which, due to the stable GFP signal, allows to follow mature OB for long periods even upon *bglap* downregulation(9). Using live-imaging approaches, we monitored the two segments bellow the amputation plane of each bony-rays and preformed a time-course analysis of OB migration every 5 hours during the first 25 hpa (Fig 1C, D). Tracking of *bglap*-positive (Bglap+) OBs revealed that OBs from the segment bellow amputation (segment 0) become motile at the 5-10 hpa time-interval and in average reach the amputation plane around 20-25 hpa (Fig 1C, D), while Bglap+ OB from the second segment (segment-1) remain predominantly immotile (Fig 1C). Subsequently, we assessed when OBs acquire proliferative capacity. Immunofluorescence for PCNA (marker of G1 phase) indicated that the proportion of Bglap+ OBs entering the cell cycle progressively increases, specifically from 12 to 24 hpa (Fig. 1E). At 24 hpa, almost 80% of Bglap+ OBs in the first segment have entered the G1 phase (Fig 1E). To examine when OBs begin to show signs of dedifferentiation towards a pre-OB state, we performed immunofluorescence for Runx2 in *bglap*:EGFP transgenic fins. In homeostasis conditions (0 hpa), Runx2 is observed in the nucleus of bone-lining mature OB (Runx2+Bglap+; Fig 1F, F’, H), which is in accordance with Runx2 being expressed at basal levels in fully differentiated OB(56). At 12 and 24 hpa we observed a slight decrease in the total number of Runx2+Bglap+ OB population within the first segment bellow amputation (Fig 1G, G’, H). OBs are not described to undergo apoptosis at this stage(8), suggesting that decrease in the Runx2+Bglap+ OBs could be due to actual loss of the *bglap* differentiation marker in a portion of dedifferentiated OB. Nevertheless, remaining Runx2+Bglap+ possess higher levels of Runx2 when compared to Runx2+Bglap+ OBs at 0 hpa (Fig 1F-G’ arrows), further indicating that they converted into a pre-OB phenotype, in accordance with published data(57). In addition, at 0 hpa a small number of pre-OB (Runx2+Bglap-), was detected in the bony-rays intersegment/joint (Fig 1F, F’ asterisk, I), which may correspond to a population of OB progenitors recently identified in this region(58). At 12 and 24 hpa, we observed an increase in the number of Runx2+Bglap- cells, suggesting that pre-OB arise in the first 12 hpa and their numbers increase until the beginning of blastema formation at 24 hpa (Fig 1F-G’, I). Altogether, these data indicate that mature OBs lose their differentiated character and contribute to the pre-OB pool in a time-window between 12 to 24 hpa. Nevertheless, we cannot exclude that pre-OB can also arise from the joint-associated OB progenitor niche. Thus, we hypothesize that at 24 hpa, multiple OB sources are recruited to contribute to the blastema (Fig 1J).

**Fig 1:**
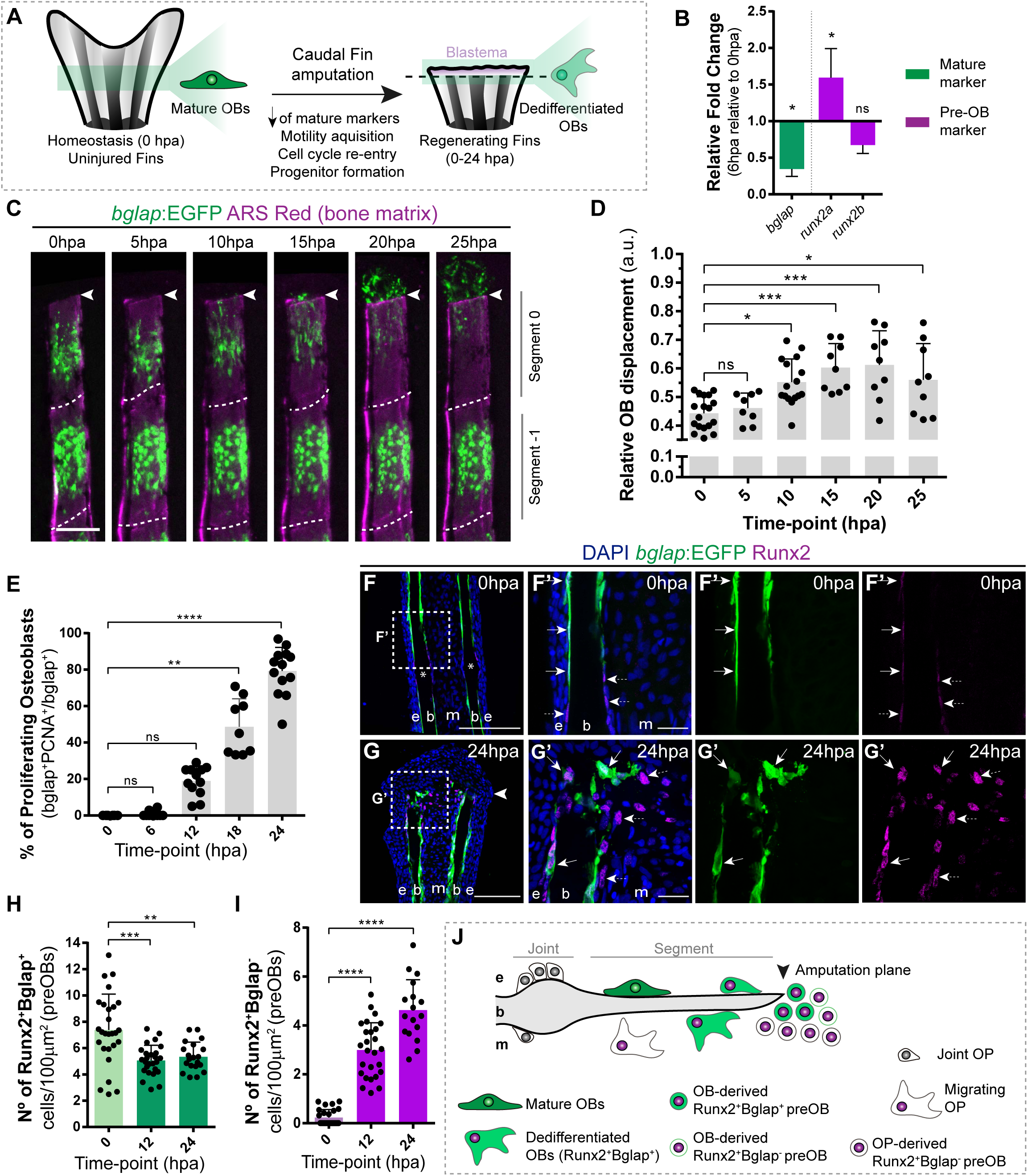
Osteoblast dedifferentiation time-window during caudal fin regeneration. (A) Biological traits of OB dedifferentiation process. (B) Relative gene expression of mature (green) and pre-OB (magenta) markers, at 6 hpa relative to 0 hpa. Statistical analysis on graph corresponds to paired t-test with Welch’s correction, Mean ± SD are displayed (n=4 biological replicates). (C) Live imaging analysis of OB motility in bglap:EGFP fish (green) during the first 25 hpa, highlighted in the segment bellow amputation (segment 0) and segment -1. Bony-rays are labelled with Alizarin red (magenta). White dashed lines delineate the intersegment region. (D) Quantification of the relative OB displacement in segment 0. Statistical analysis displayed on graph corresponds to Kruskal-Wallis test with Mean ± SD (n=9-18 bony-rays). (E) Percentage of proliferating OBs through immunofluorescence against PCNA in bglap:EGFP fish. Statistical analysis displayed on graph corresponds to Kruskal-Wallis test with Mean ± SD (n=9-13 cryosections). (F-G’) Representative cryosection images of bglap:EGFP (green) fins immunostained for Runx2 (magenta) and counterstained with DAPI (blue), in (F, F’) uninjured fish and (G, G’) at 24 hpa; arrows indicate Runx2+Bglap+ cells and dashed arrows indicate Runx2+Bglap- cells. (H, I) Quantification of (H) Runx2+Bglap+ and (I) Runx2+Bglap- cells during the first 24hpa. Statistical analysis displayed on graph corresponds to Mann-Whitney test with Mean ± SD (n= 18-27 cryosections). (J) Cellular sources that contribute for new pre-OBs formation after injury include mature osteoblasts and potentially joint OP. White arrowhead indicates amputation plane and dashed squares represent magnified panels in F’ and G’. E: epidermis; b: bone; m: mesenchyme; ns: not significative; *P <0.05; **P<0,01; ***P<0,001; ****P<0,0001. Scale bars represent 100 µm and 30 µm in magnified panels.

Here we show that mature OB dedifferentiation is an early response to injury and entails important transcriptional and phenotypic alterations, occurring in a narrow time-window between 6-12 hpa, before blastema induction and during the wound healing phase.

### 2. Metabolic reprogramming towards glycolysis occurs at early stages of fin regeneration prior to blastema formation

Having established that the first hours after caudal fin amputation are crucial for the activation of OBs, we proceeded with the identification of the initial regulators of OB dedifferentiation. We isolated OBs from the first bony-ray segment bellow amputation at 0 hpa, our control population and closest to a homeostatic state, and at 6 hpa, when most dedifferentiation features are not detected yet, and performed transcriptome analysis (Sup Fig 1A). OBs were isolated by FACS using the *bglap*:EGFP transgenic line (Sup Fig 1B). We then compared the expression profiles of mature OBs (0 hpa) and dedifferentiating OBs (6 hpa) and performed a gene ontology analysis to evaluate whether our set of differentially expressed genes is associated with a specific biological process or signalling pathway, particularly relevant for OB dedifferentiation. We observed that OBs show a dynamic transcriptional response at 6 hpa in comparison to OBs from uninjured conditions with almost 2200 differentially expressed genes, from which 1040 were downregulated and 1130 were upregulated (Sup Fig 1C, D). These data further demonstrating that the dedifferentiation machinery is triggered very early during regeneration. Importantly, a large set of genes related to energy metabolism was also dramatically altered (Sup Fig 1C, D).

Depending on the energy and biomass demands, cells can uptake glucose and, through glycolysis, use it to produce pyruvate, which can serve as a substrate to: generate acetyl-CoA and fuel mitochondrial OXPHOS, obtaining a high energy yield; or to produce lactate, allowing diversion of metabolic intermediates from glycolysis towards various biosynthetic pathways(59, 60) (Fig 2A). We observed that at 6 hpa OBs upregulate the expression of major glycolytic enzymes, such as *hexokinase 1* (*hk1*) and *phosphofructokinase* (*pfkp*), whereas OXPHOS components remain mostly unchanged (Fig 2B). Most importantly, we detected a significant increase in *lactate dehydrogenase* (*ldha*) expression, indicating an increase in lactate production, thereby diverting pyruvate from mitochondrial oxidation. (Fig 2B). These data suggest that dedifferentiating OBs change their metabolic signature, adopting a metabolic program that prioritises glycolysis. Since OBs only account for around 1-3% of the total cell number in a bony-ray segment (Sup Fig 1B), we evaluated whether these changes in gene expression were specific to OB undergoing dedifferentiation or are also observed as a general behaviour induced upon injury. For that, we collected the first segment below the amputation and analysed by qPCR the expression profile of key glycolytic and OXPHOS genes in regenerating caudal fins from 6 hpa and 24 hpa and compared to control caudal fins (0 hpa). At 6 hpa, when cells start to dedifferentiate, most glycolytic enzymes were highly upregulated, including *hk1*, *hk*2, *pfkpa*, and *pyruvate kinase* (*pkma)* (Fig 2C). Importantly, as observed during OB dedifferentiation, *ldha* expression was increased and *pyruvate dehydrogenase a 1b* (*pdha1b*) was downregulated. *pdha1b* is part of the Pdh1 complex, which catalyses irreversibly the conversion of pyruvate to acetyl-CoA and progression to OXPHOS, further corroborating that at this stage pyruvate is being shunt from the mitochondria and stimulating lactate-producing glycolysis (Fig 2C). As for the OXPHOS components analysed, most remained unchanged or slightly upregulated. However, this increase in expression was not as accentuated as the one observed for the glycolytic pathway (Fig 2C). Later, at 24 hpa, when most cells have dedifferentiated and the blastema starts to be assembled, we observed that most glycolytic enzymes were still significantly upregulated, although not as striking as at 6 hpa time-point, and *ldha* expression continued to be upregulated (Fig 2D). At this time-point, we also observed an increase in expression of some OXPHOS components (Fig 2D), suggesting that later stages of blastema formation may also require mitochondrial glucose oxidation. Consistent upregulation of the genes associated with glycolysis and lactate production, suggests that the OB dedifferentiation process and possibly the early response to amputation is characterized by a metabolic reprogramming towards non-oxidative glycolysis.

**Fig 2.**
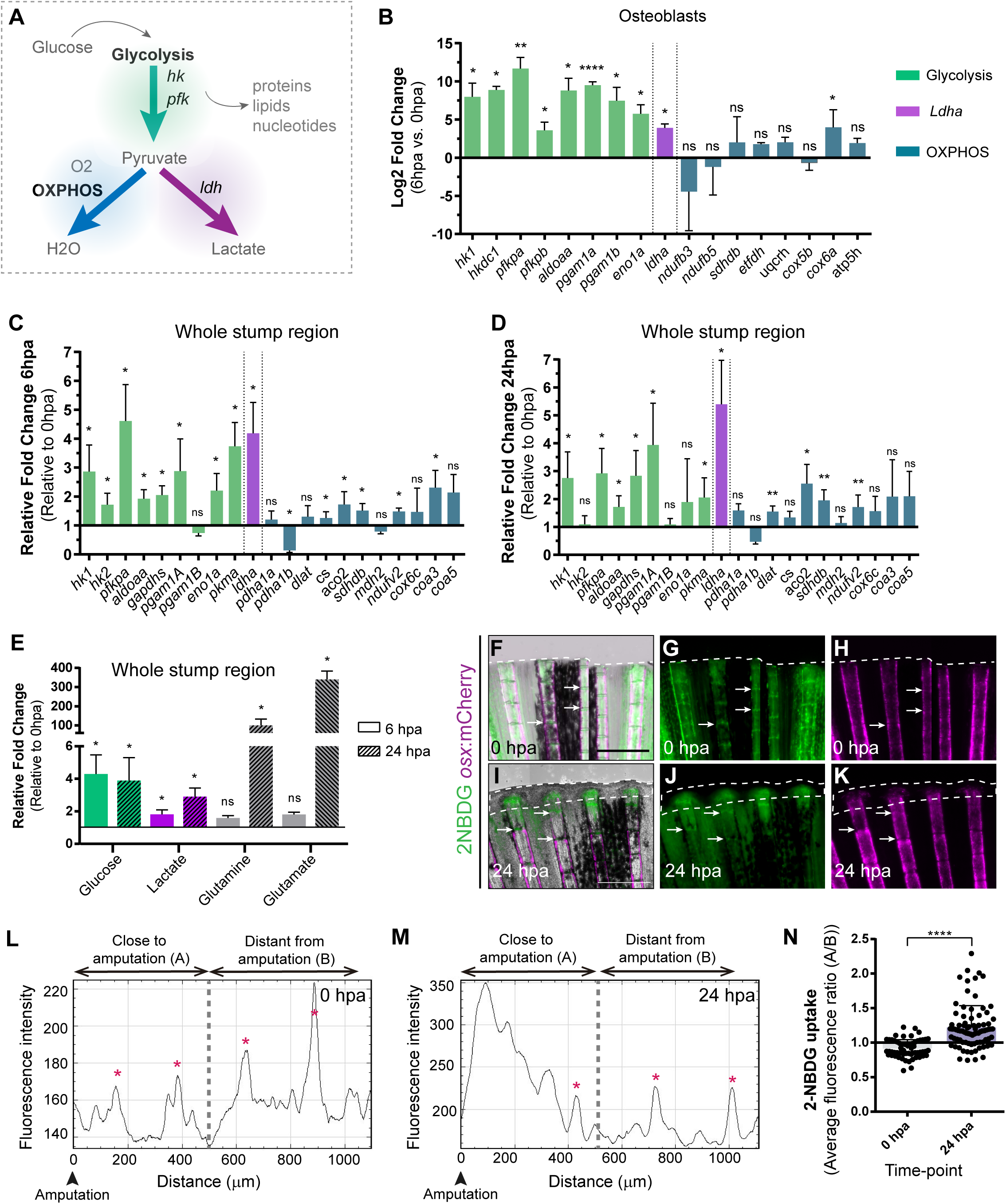
Metabolic reprogramming is triggered during zebrafish caudal fin regeneration. (A) Schematic representation of glucose metabolism. (B) OB gene expression profile of glycolytic enzymes (green), *ldha* (magenta) and OXPHOS components (blue) at 6 hpa relative to uninjured conditions (0 hpa). Statistical analysis with t test and Welch’s correction (n=3 biological replicates), Mean ± SD are displayed. (C, D) Relative gene expression of glycolytic enzymes (green), *ldha* (magenta) and OXPHOS components (blue), in the whole caudal fin stump, at (C) 6 hpa and at (D) 24 hpa in comparison to uninjured conditions (0 hpa). Statistical analysis with paired t test (n=5 (C) and 4 (D) biological replicates). (E) Metabolite measurements at 6 hpa (clean columns) and 24 hpa (stroked columns) in relation to uninjured conditions (0 hpa), in the whole caudal fin stump. Statistical analysis with Mann-Whitney test (n=4 biological replicates). (F-K) Live imaging of 2NDBG uptake (green) in osx:mCherry caudal fins (magenta) at (F-H) 0 hpa and (I-K) 24 hpa. Arrows indicate uptake of 2NBDG in the intersegment regions. White dashed line delineates the regenerated tissue. Scale bar represents 500 µm. (L-M) Intensity of 2NBDG uptake in regions close and distant to the amputation site, at (L) 0 hpa and (M) 24 hpa. Red * indicate peaks of 2NDBG uptake in the intersegments. (N) Ratio of 2NBDG uptake at 0 hpa and 24 hpa. Statistical analysis on graph corresponds to Mann-Whitney test, Mean ± SD are displayed (n=54 and 83 bony-rays). ns: not significative; *P <0.05; **P<0,01; ****P<0,0001.

Nevertheless, gene expression data may not reflect an actual activation of a specific metabolic pathway. Therefore, to corroborate these observations, we used mass spectrometry (MS) to quantify specific metabolites and determine the prevalent energy metabolism route. For that, we dissected the whole fin stump, and analysed the amount of glucose, lactate, glutamine and glutamate at 0, 6 and 24 hpa. According with our transcriptome data, we observed an increase in glucose and lactate at 6 and 24 hpa in relation to control fins (0 hpa), which is consistent with an increment in the glycolytic influx (Fig 2E). We also observed a strong increase of glutamine and glutamate at 24 hpa but not at 6 hpa (Fig 2E). Glutamine and glutamate act as important substrates for protein and nucleotide synthesis necessary to support cellular integrity and growth suggesting an increase in biosynthesis(61, 62).

We also monitored glucose uptake during regeneration using a fluorescent glucose analogue (2- NBDG). In control fins (0 hpa), we detected glucose uptake in the intersegment region (Fig 2F-G arrows, L, N), where a population of OP have been identified(58), which may imply that under homeostatic conditions, these cells require higher levels of glucose to maintain the progenitor niche. At 24 hpa we observed a significant increase of glucose uptake in the blastema primordium and in the first segment bellow the amputation (Fig 2I-K, M, N).

Thus, metabolic reprograming seems to constitute an integral response to amputation, tightly regulated in terms of time and space. Our transcriptome and metabolome studies indicate a specific metabolic signature as part of an early response to amputation by OBs and by the remaining caudal fin tissue. Overall, cells respond to injury by increasing the glycolytic influx and lactate production just preceding dedifferentiation and blastema formation. Corroborating the hypothesis that amputation induces a metabolic reprogramming in the form of a glycolytic switch, which may be part of a general program that coordinates the regenerative response.

### 3. Glycolytic influx and lactate generation supports blastema formation during caudal fin regeneration

To investigate the functional relevance of the energy metabolism, we chose to start with a broader analysis in which we manipulated specific branches of energy metabolism with used pharmacological compounds and address its requirement for blastema formation. The blastema is hallmark and prerequisite of epimorphic regeneration (Fig 3A) and is fully assembled within 48 hpa(4–6). Firstly, we inhibited glycolysis using an established glucose analogue, 2-Deoxy-D-glucose (2DG)(52, 53), that is non-metabolized by Hk, the rate limiting step of glycolysis. 2DG was administered in different time points right after amputation to affect different stages of blastema formation: for a short period before blastema formation (0 hpa and 0-12hpa), or for a prolonged period, which includes the blastema assembly phase (0-24hpa and 0-36hpa). Fins were imaged at 48 hpa to determine the effect of 2DG on blastema growth by measuring the total fin regenerated area (Fig 3B). We observed that the higher the duration of the treatment the stronger the effect on blastema growth in comparison to control fins (Fig 3C, D-K). Interestingly, a single dose of 2DG administered at 0 hpa already showed significant smaller regenerates when compared to control fins (Fig 3C-E). Administration of 2DG between 0-12, 0-24 and 0-36 hpa had a dose-dependent inhibitory effect on the overall regenerated area (Fig 3C, F-G). Accordingly, blocking of the glycolytic influx for the first 36 hpa led to a complete abrogation of blastema assembly (Fig 3C, J-K). Similarly, the glycolysis inhibitor acting downstream of 2DG, 3-(3-pyridinyl)-1-(4-pyridinyl)-2-propen-1-one (3PO), which partially inhibits the glycolytic activator of Pfk (63) (Fig 3A), also exhibited a general impairment of caudal fin regeneration, as observed at 48 hpa in relation to control fins(Sup Fig 2A-D).

**Fig 3.**
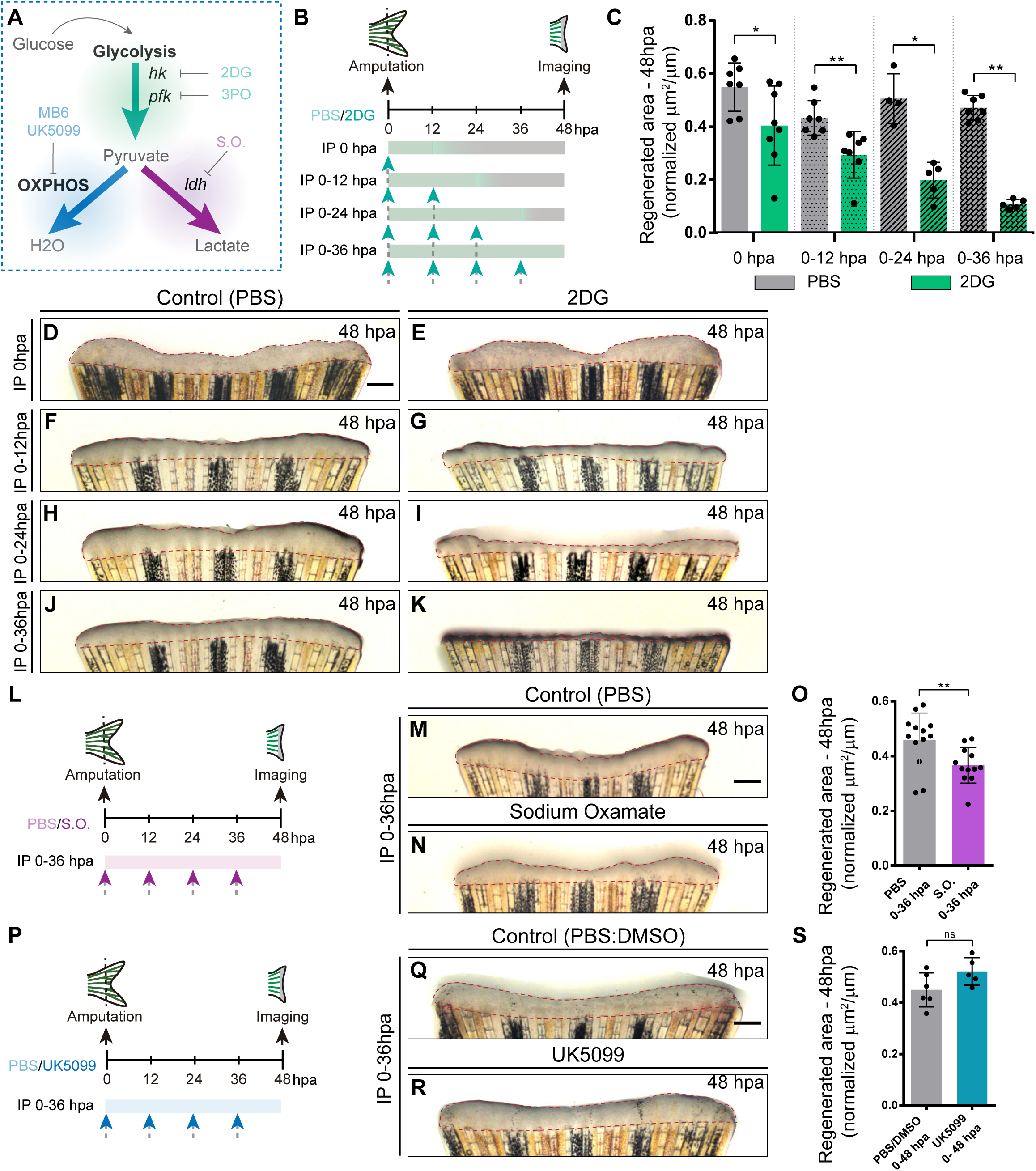
Inhibition of glycolysis, but not OXPHOS, impairs blastema formation. (A) Schematic representation of the compounds used to manipulate glucose metabolism. (B) Experimental design used to inhibit the glycolytic influx during fin regeneration. Control and treated fish are administered, via IP injection, with vehicle (PBS) or glycolytic inhibitor, 2DG, respectively, every 12 hours, from fin amputation (0 hpa) until 48 hpa. Different time-intervals were used for injections: 0 hpa) IP injection at 0 hpa; 0-12 hpa) IP injection at 0 and 12 hpa; 0-24 hpa) IP injection at 0, 12 and 24 hpa; 0-36 hpa) IP injection at 0, 12, 24 and 36 hpa. (C) Quantification of the total fin regenerated area at 48 hpa, after vehicle (PBS) or 2DG injection, at specific time-intervals during regeneration. (D-K) Representative images of 48 hpa fins treated with (D,F,H,J) vehicle (PBS) or (E,G,I,K) 2DG during different time-intervals. (L) Experimental design used to inhibit the lactate formation during fin regeneration. Fish are administered, via IP injection, with vehicle (PBS) or S.O. every 12 hours, from fin amputation (0 hpa) until 48 hpa. (M, N) Representative images of 48 hpa CF treated with (M) vehicle (PBS) or (N) S.O.. (O) Quantification of the total fin regenerated area at 48 hpa, after vehicle (PBS) or with S.O. injection. (P) Experimental design used to inhibit pyruvate translocation to mitochondria during fin regeneration. Fish are administered, via IP injection, with vehicle (PBS) or MPC inhibitor, UK5099, every 12 hours from fin amputation (0 hpa) until 48 hpa. (Q, R) Representative images of 48 hpa fins treated with (Q) PBS (control) or (R) UK5099. (S) Quantification of the total fin regenerated area at 48 hpa, after vehicle (PBS) or with UK5099 injection. Statistical analysis displayed on each graph corresponds to Mann-Whitney test with Mean ± SD, each dot corresponds to one fish. Scale bar represents 500 µm. Dashed lines define the regenerated tissue. * p< 0.05, ** p< 0.01.

Next, we blocked the conversion of pyruvate to lactate using the inhibitor of Ldh, Sodium Oxamate (S.O.)(64) (Fig 3A). S.O. administration during the first 36 hpa resulted in a decrease of the caudal fin regenerated area in respect to control caudal fins (Fig 3L-O), although exhibiting a milder effect when compared to glycolysis inhibition (Fig 3C-K and Sup Fig 2A-D). In contrast, inhibition of OXPHOS using two independent compounds, a mitochondrial pyruvate carrier inhibitor (UK5099)(65) and an inhibitor of the mitochondrial electron transport chain (MitoBlock-6 (MB6))(66), had no effect on total regenerated area and thus on blastema formation at 48 hpa when compared to control fins (Fig 3P-S and Sup Fig 2E-H). Together, these results indicate that glycolysis and lactate generation, but not OXPHOS, are essential for the initial stages of caudal fin regeneration. Our data strongly supports the hypothesis that the cells that respond to amputation reprogram their metabolic profile since very early thereby enhancing the glycolytic machinery/influx to support blastema formation.

### 4. Inhibition of glycolysis interferes with osteoblast dedifferentiation and pre-osteoblast pool assembly and proliferation

Considering that glycolysis inhibition culminated in a severe impairment of blastema assembly, we investigated whether those defects could be due to a role of glycolysis in mediating cell dedifferentiation and/or re-acquisition of proliferative capacity (Fig 4A), processes that precede and are indispensable for blastema formation. Therefore, we blocked glycolysis with 2DG and performed a detailed characterization of OB dedifferentiation at 24 hpa (Fig 4B). We began by analysing the expression profile of OB markers that serve as a read-out of their dedifferentiated status, namely *bglap* and *runx2*. We observed that inhibition of the glycolytic influx led to a significant upregulation of *bglap* and to a decrease in both *runx2* orthologues (Fig 4C), as opposed to what happens in OB undergoing dedifferentiation in normal regenerating conditions(8, 9). Immunofluorescence analysis for Runx2 in *bglap*:EGFP reporter fish, showed that 2DG treatment had no effect in the number of dedifferentiated Runx2+Bglap+ pre-OBs (Fig 4E-G). However, the number of Runx2+Bglap- pre-OBs was significantly reduced (Fig 4E-G) when compared to control fins (Fig 4D, F, G). Overall, these data indicate that blocking glycolytic influx hampers mature OB dedifferentiation, which become unable to operate as a source of pre-OB resulting in impaired pre-OB pool assembly within the blastema primordium.

**Fig 4.**
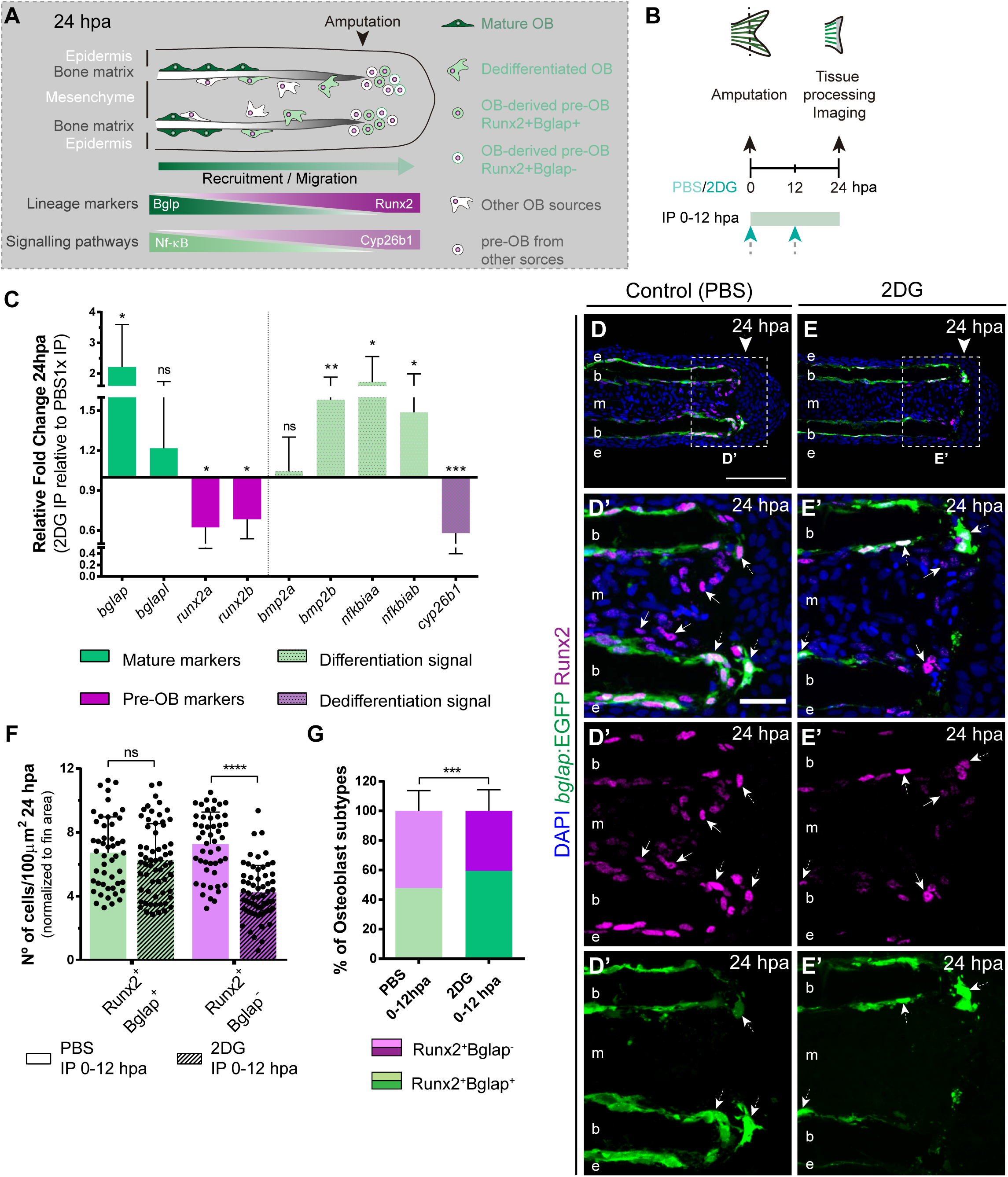
Inhibition of glycolysis impairs osteoblast dedifferentiation. (A) Schematic representation of pre-OBs formation during regeneration. Pre-OBs arise from OB dedifferentiation and potentially from the joint OP niche. OB dedifferentiation is correlated with inactivation of NF-ΚB and increase in Cyp26b1 activity. (B) Experimental design used to inhibit glycolysis. Fish are administered, via IP injection, with vehicle (PBS) or 2DG, from fin amputation (0 hpa) until 24 hpa. (C) Relative gene expression of mature and pre-OBs markers, and differentiation and dedifferentiation pathways, in the whole fin stump at 24hpa, in 2DG treated fins compared to control condition (0 hpa). Statistical analysis with paired t-test (n=8 biological replicates). (D-E’) Representative cryosection images of 24 hpa bglap:EGFP (green) caudal fins immunostained for Runx2 (magenta) and counterstained with DAPI (blue), in fish treated with (D,D’) vehicle (PBS) or (E,E’) 2DG. White dashed boxes delineate magnified panels in D’ and E’. Arrows indicate Runx2+Bglap- cells. Dashed arrows indicate Runx2+Bglap+ cells. Arrowhead indicates amputation plane. E: epidermis; b: bone; m: mesenchyme. Scale bar represents 100 µm and 30 µm in magnified panels. (F) Total number of Runx2+Bglap+ and Runx2+Bglap- cells per area at 24 hpa. (G) Percentage of Runx2+Bglap+ and Runx2+Bglap- OBs subtypes. Statistical analysis displayed on each graph corresponds to Mann-Whitney test with Mean ± SD (n=49 (PBS) and 69 (2DG) cryosections). ns: not significant;** p< 0,01; *** p<0,001; ****p<0,0001.

To further validate our observations, we decided to evaluate whether the pathways proposed to mediate mature OB dedifferentiation where also altered upon glycolysis inhibition. During homeostasis NF-kB pathway maintains retinoic acid (RA) signalling in OBs, supporting differentiation. Upon amputation, NF-kB pathway becomes inactivated and RA is degraded through activity of the RA-degrading enzyme, Cyp26b1, thereby inducing OB dedifferentiation(57, 67) (Fig 4A). In addition, Bmp signalling is also considered to be a potent inducer of OB differentiation during regeneration(12,13,68,69). Thus, we performed qPCR analysis at 24 hpa of NF-kB target genes (*e.g. nf*k*biaa* and *nf*k*biab*), retinoic acid degrading enzyme (*e.g. cyp26b1*) and Bmp ligands (*e.g. bmp2a* and *bmp2b*) in control and 2DG-treated fins (Fig 4C). We observed an increase in NF-kB target genes and in *bmp2b*, accompanied by a decrease in *cyp26b1* in 2DG-treated fins. Thus, suppression of glycolysis seems to maintain the mature OB differentiation profile and to prevent their dedifferentiation after amputation, resembling a pre-amputation scenario.

We then evaluated whether glycolysis is necessary for other aspects all aspects of OB dedifferentiation namely migration towards the stump and cell cycle re-entry. Surprisingly, by measuring the relative cell displacement of the OB population in the first segment bellow the amputation region after 24 hpa, we observed that 2DG administration had no effect on the ability of OB to migrate and reach the amputation zone (Fig 5A-E). However, cell cycle re-entry, examined through a EdU 3-h pulse assay in *bglap*:EGFP transgenics, was strongly affected by 2DG treatment (Fig 5F-G’) contrasting with control caudal fins. We observed a significant reduction of total number of Runx2+Bglap+EdU+ pre-OBs per area (Fig 5H) and of percentage of Runx2+Bglap+EdU+ within the Runx2+Bglap+ population (Fig 5I). By quantifying the remaining pre-OB population Runx2+Bglap-, we also observed a reduction in the number and relative percentage of Runx2+Bglap-EdU+ pre-OBs upon 2DG treatment (Fig 5F-G’ arrows, H. I). This indicates that blocking glycolytic machinery has a severe impact in the number of pre-OB that re-enter the cell cycle, reducing their proliferative capacity. This inhibition of cell cycle re-entry by 2DG was also observed in the blastema primordium mesenchyme (Sup Fig 3A-F, H) and the overlying epidermal cap (Sup Fig 3A-F, G). Moreover, the decline in the capacity of pre-OB to re-entre the cell cycle and to proliferate could be correlated with defects in the activity of pathways known to be indispensable for blastema proliferation, such as, Wnt, Insulin and Fgf signalling pathways(70–75). This is predicted based on our results showing a downregulation of *wnt10a*, *igf2b* and *fgf20a* in 2DG- treated caudal fins in comparison to controls (Fig 5J). We further complement and confirmed our results using 3PO, a partial glycolytic inhibitor, in *bglap*:EGFP transgenic fish. In accordance with previous results 3PO led to a reduction in cell cycle re-entry of Bglap+ pre-OB, mesenchymal and epidermal cells, measured by PCNA immunofluorescence (Sup Fig 3I-O).

**Fig 5.**
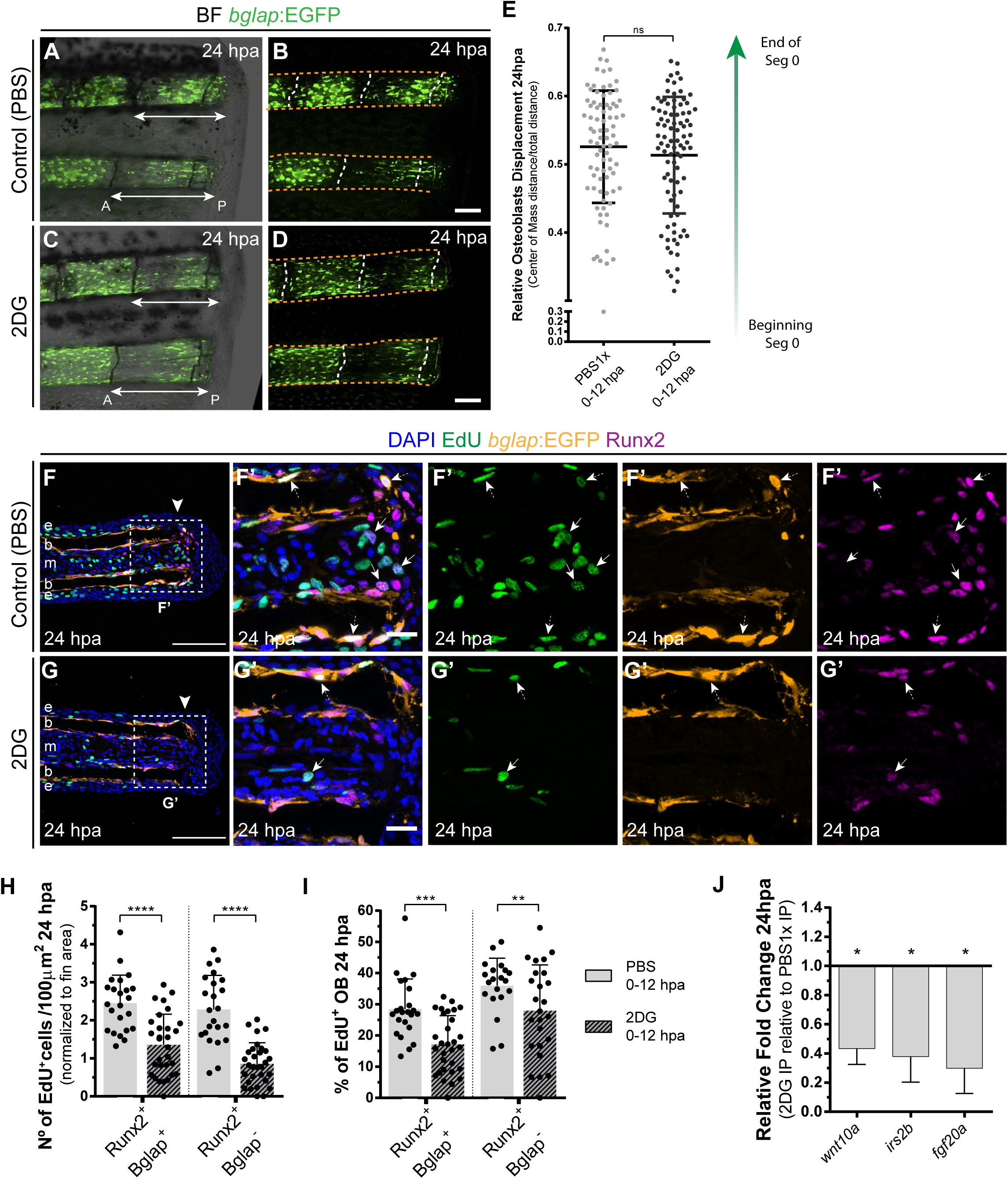
Inhibition of glycolysis impairs osteoblast cell cycle-entry. (A-D) Representative images of bglap:EGFP caudal fins at 24 hpa, treated with (A-B) vehicle (PBS) or (C-D) 2DG. Double white arrows indicate the anterior (A) and posterior (P) axis. White dashed lines indicate intersegment regions. Orange dashes lines delineate the bony-ray surface. (E) Measurement of relative OB displacement along segment 0, below the amputation plane, at 24 hpa in fins treated with vehicle (PBS) or 2DG. (F-G’) Representative cryosection images of 24 hpa bglap:EGFP (orange) caudal fins immunostained for Runx2 (magenta), labeled with EdU (green) and counterstained with DAPI (blue), in fish treated with (F) control (PBS) or (G) 2DG. Dashed boxes delineate amplified panels in F’ and G’. Arrows indicate proliferative EdU+ Runx2+Bglap- cells. Dashed arrows indicate proliferative EdU+ Runx2+Bglap+ cells. Arrowhead indicates amputation plane. Scale bar represents 100 µm and 30 µm in amplified panels. (H) Total number of Runx2+Bglap+ and Runx2+Bglap- cells at 24hpa, in fins treated with vehicle (PBS) or 2DG. (I) Percentage of proliferative Runx2+Bglap+ and Runx2+Bglap- cells at 24hpa, in fins treated with vehicle (PBS) or 2DG. Statistical analysis displayed on each graph corresponds to Mann-Whitney test with Mean ± SD (n=23-30 cryosections). (J) Relative gene expression at 24 hpa in 2DG treated fins, compared to control. Statistical analysis with unpaired t test and Welch’s correction (n=5 biological replicates). ns: not significant; *P <0,05; **P <0,01; ***P<0,001; **** P<0,0001.

To exclude any effect of glycolysis inhibition on cell survival that could interfere with our observations, we performed a TUNEL assay at 24 hpa in controls and in 2DG-treated *bglap*:EGFP transgenic zebrafish. Except for the mesenchyme compartment, which showed a slight increase in the number of TUNEL-positive cells, the epidermis and Runx2+ pre-OBs showed no major alterations upon 2DG treatment (Sup Fig 4A-L). This suggest that glycolysis may support cell survival in the mesenchymal compartment, but not in the epidermis or in the pre-OB pools. Thus, blocking glycolysis is sufficient to inhibit OB dedifferentiation and cell cycle re-entry, without affecting their capacity to migrate or survive.

Taken together our results reveal that enhancing glycolysis induces mature OB plasticity by promoting their dedifferentiation into pre-OB and enabling pre-OB and other lineages in the to re-acquire proliferative capacity within the blastema. We also provide evidence that energy metabolism controls these aspects of OB response to injury through a glycolysis-dependent transcriptional regulation. Therefore, these results provide solid evidence that metabolic reprogramming towards glycolysis is a novel and powerful conductor of the cell fate changes and cell cycle re-entry preceding blastema assembly.

### 5. Glycolysis suppression leads to impaired blastema proliferation and defective distribution of new osteoblast subtypes

Based on our results so far, we demonstrated a fundamental role of glycolysis in governing regulating OB dedifferentiation and the early stages of blastema formation. Subsequently, we aimed to investigate how prolonged inhibition of glycolysis would interfere with blastema organization and with *de novo* OB formation (Fig 6A). At 48 hpa, the blastema is subdivided in a patterning zone (PZ) and in a proximal (PB) and distal (DB) compartments. These regions are characterized by distinct OB subtypes, based on their maturation and proliferative state, exhibiting a proximal-distal hierarchical distribution (70,76–78): show cycling Runx2^+^ pre-OBs maintain the progenitor pool in the DB and as they proliferate and populate the PB, they differentiate into fast-proliferating Runx2^+^ Osx^+^ immature OBs (12, 14), which will later give rise to non-proliferative differentiated OBs in the PZ (Fig 6B)(12, 14). To determine how glycolysis affects the general distribution of OB subtypes within the blastema, we exposed *runx2*:EGFP and *osx*:mCherry zebrafish to 2DG treatment thought the first 48 hpa (Fig 6A). We observed that prolonged 2DG-treatment caused a severe abrogation of blastema organization (Sup Fig 5A-H) with a strong reduction in *osx* (Sup Fig 5B, F, D, H) and *runx2* (Sup Fig 5C, D, G, H) when compared to control fins. Indicating that 2DG administration strongly alters OB specific gene expression within the blastema. In normal regenerating conditions, immunofluorescence for Runx2 in *osx*:mCherry transgenics display a proper organization of the OB subtypes within the blastema (Fig 6C- C’). The cluster of Runx2+ pre-OBs, referred as the Runx2+Osx- subtype, is restricted to the DB and represents only a small fraction of the OB lineage in the blastema (Fig 6C, C’, E), while immature OBs, referred as the Runx2+Osx- subtype, reside in the PB region and correspond to the major OB subtype (Fig 6C, C’, F). In contrast, we observe that 2DG-treated fish had an accentuated decrease in the total number of Runx2^+^Osx^+^ immature OBs (Fig 6D, D’, F), while distal Runx2^+^Osx^-^ pre-OBs remain unchanged (Fig 6D, D’, E), leading to an imbalance between the OB populations within the blastema (Fig 6G). Similar results were obtained when using the glycolytic inhibitor 3PO (Sup Fig 5I-L).

**Fig 6.**
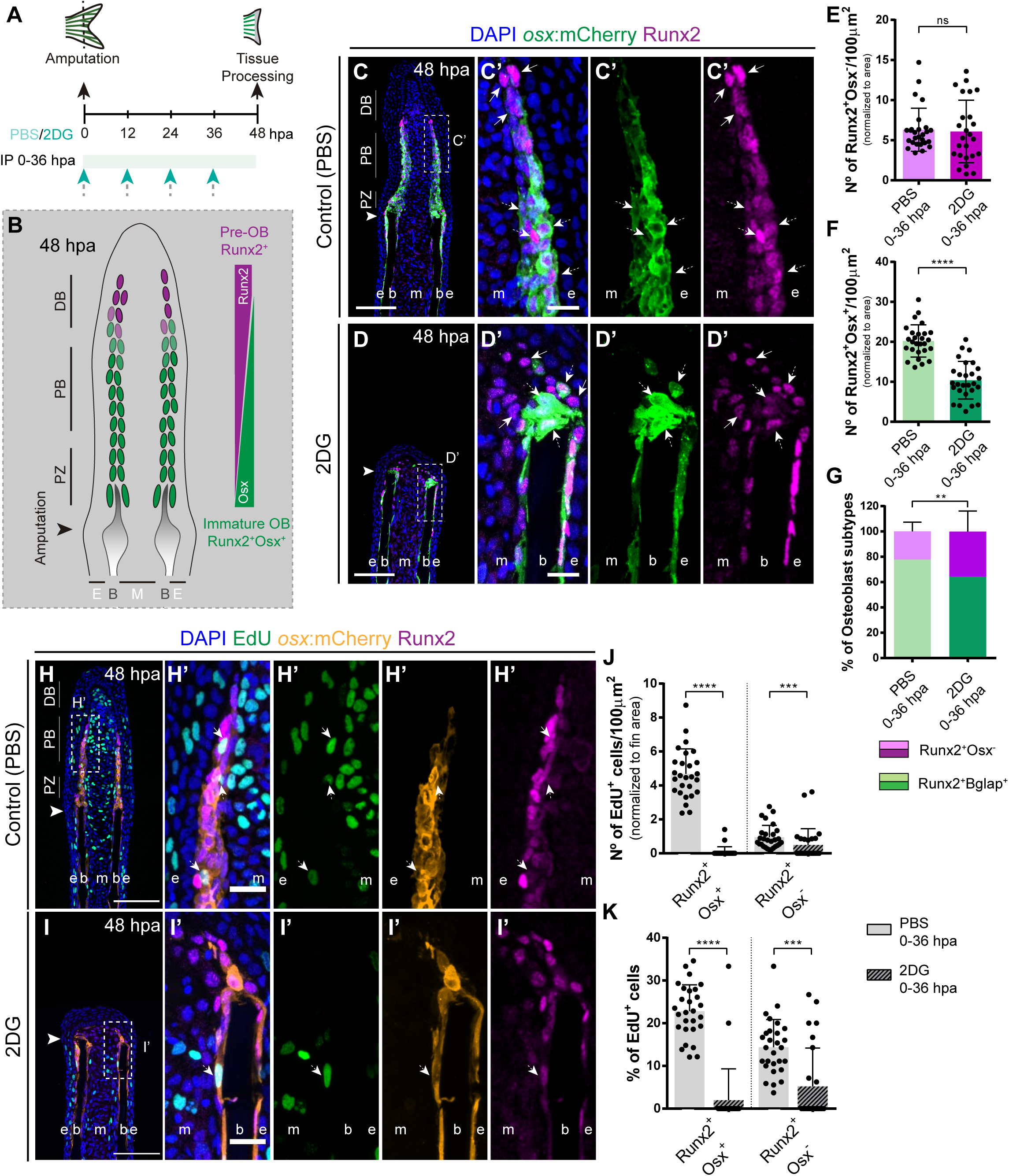
Inhibition of glycolysis affects formation of osteoblast subtypes and proliferation within the blastema. (A) Experimental design used to inhibit glycolysis. Fish are administered, via IP injection, with control (PBS) or 2DG every 12 hours, from fin amputation (0 hpa) until 48 hpa. (B) Schematic representation of the distribution of OBs subtypes along the blastema. (C-D) Representative cryosection images of 48 hpa osx:mCherry (green) caudal fins immunostained for Runx2 (magenta) and counterstained with DAPI (blue), in fish treated with (C,C’) PBS and (D,D’) 2DG. Dashed boxes represent magnified panels in C’ and D’. Arrows indicate Runx2+ Osx- pre-OBs. Dashed arrows indicate Runx2+ Osx+ immature OBs. (E-F) Total number of (E) Run2+Osx- and (F) Runx2+Osx+ subtypes in 48 hpa fins treated with PBS or 2DG. (G) Percentage of Runx2+/Osx- and Runx2+Osx+ subtypes in 48 hpa fins treated with PBS or 2DG. (H-I’) Representative cryosection images of 48 hpa osx:mCherry (orange) caudal fins immunostained for Runx2 (magenta), EdU (green) and counterstained with DAPI (blue), in fish treated with (H,H’) PBS and (I,I’) 2DG. Dashed boxes represent magnified panels in H’ and I’. Arrows indicate proliferative Edu+ Runx2+ Osx- pre-OBs. Dashed arrows indicate proliferative Edu+ Runx2+ Osx+ immature OBs. Arrowheads indicate amputation plane. Scale bar represents 100 µm and 20 µm in magnified panels. (J) Total number of Runx2+Osx+ and Runx2+Osx- proliferative OBs subtypes at 48 hpa fins, treated with PBS or 2DG. (K) Percentage of proliferative Runx2+Osx+ and Runx2+Osx- OBs subtypes in 48 hpa CFs, treated with PBS or 2DG. DB: distal blastema; PB: proximal blastema; PZ: patterning zone; E: epidermis; B: bone; M: mesenchyme. ns: not significant; **P <0,01; ***P<0,001; **** P<0,0001.

Afterwards, we decided to ascertain if the proliferative abilities of each OB subtype in control and 2DG-treated fish, through a EdU 3 h-pulse assay. As previously reported(12), in control fins Runx2+Osx+ immature OBs exhibit a higher proliferation rate than Runx2+Osx- pre-OBs (Fig 6H-K). Strikingly, glycolysis inhibition had a profound impact on proliferation at 48 hpa (Fig 6I-K) than at 24 hpa (Fig 5F-I), with both OB subtypes exhibiting a significant reduction of their proliferative capacity (Fig 6H-K). This reduction was particularly noticeable in Runx2+Osx+ OBs, which are fast-proliferating cells in control conditions. We also noticed that, as observed at 24 hpa (Sup Fig 3G, H), the epidermis and mesenchyme were also affected, displaying a decrease in proliferation in 2-DG-treated fish (Sup Fig 6A-H). Given the results obtained, the most likely interpretation is that both pre-OB and immature OB populations accumulate at the stump region since they are unable to proceed in the cell cycle and divide. It is noteworthy mentioning that after 2DG administration, Runx2+Osx- pre-OBs are still able to be recruited at this stage, but do not increase in total numbers due to inability to self-renew. Interestingly, this may indicate that, although OB dedifferentiation is compromised after blocking the glycolytic influx, pre-OB may be generated by alternative sources.

Overall, these data demonstrate an indispensable role of glycolysis in regulating blastema proliferation and compartmentalization with important implications for new OB generation.

## Discussion

### Metabolic reprogramming as an early response to caudal fin injury

OB dedifferentiation has been suggested to occur at the end of wound healing phase (0-18 hpa) and during the blastema induction phase (12-24 hpa)(8,9,12). Here, by providing a deeper characterization of OB dedifferentiation, we demonstrate that this process is triggered as early as 6 hpa, in parallel with the initial wound healing response(79). Moreover, our transcriptomic analysis of isolated OBs revealed a dynamic transcriptional response at 6 hpa in comparison to OBs from uninjured conditions. This provides the first molecular characterization of OBs preceding the dedifferentiation stage, highlighting that mature OBs start changing their transcriptome earlier than expected and that the first after amputation are crucial for the transcriptional and phenotypic alterations leading to dedifferentiation. The set of differentially expressed genes unveils potential new players worth revisiting in the future. Our study uncouples OB response from surrounding tissues, and addresses the early stages of fin regeneration, which are the least investigated. In fact, most published data focus on time points from 24 hpa onwards, when wound closure has finished, blastema formation is in progress and consequently initial cell identity transitions have been dictated, potentially missing initial regulators of dedifferentiation. Importantly, we show that at 6 hpa OB prioritise lactate-producing glycolysis over OXPHOS, when compared to OB from uninjured fins. Additionally, we also observed a similar response at 6 hpa in the whole fin stump, corroborated by gene expression and metabolomic data. These alterations persist at least until 24 hpa, when the blastema primordium is being assembled. We also show that glycolysis is indispensable to support blastema formation and regeneration. Blocking glycolysis leads to a complete blastema suppression, and even a single injection of 2DG at 0 hpa was sufficient to induce aberrant blastema formation. These results indicate that OBs and other cell lineages respond to amputation by undergoing a metabolic reprogramming that favours glycolysis over OXPHOS (Fig 7). Furthermore, glycolysis is necessary from the early onset of regeneration and that the time interval when these changes in metabolism happen appears to be fundamental for the initiation of regeneration. It is possible that early wound response signals are important to trigger the metabolic shift. One potentially relevant event described at this stage is reactive oxygen species (ROS) production(80). ROS and cellular metabolism are tightly connected as ROS are a by-product of mitochondrial oxidation(81) and produced by epithelial cells upon damage (82). ROS are shown to activate important molecules, such as HIF-1α, which has been shown to promote metabolic reprogramming towards glycolysis in other contexts(83–85). It would be interesting to evaluate whether ROS production is necessary to induce metabolic reprogramming during caudal fin regeneration.

**Fig 7.**
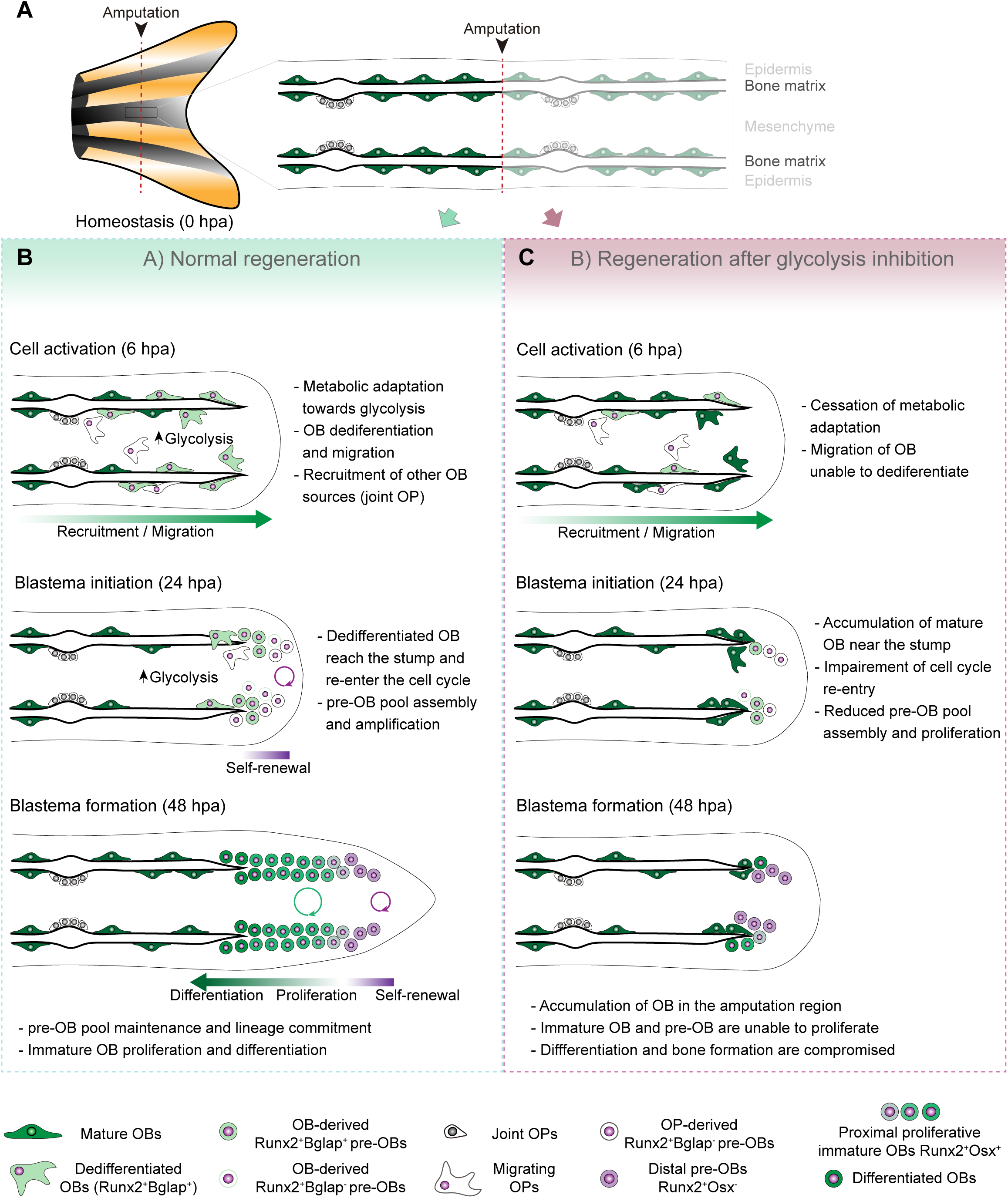
Model for the role of metabolic reprogramming during caudal fin regeneration. (A) In homeostasis, mature OBs reside in close contact with the bony-ray surface, secreting the collagenous bone matrix. Upon caudal fin amputation, OBs and other cell types in the regenerating fin respond by undergoing a metabolic reprogramming in the shape of a glycolysis shift that is essential for regeneration to proceed. Enhancing glycolysis promotes OB dedifferentiation, by releasing Cyp26b1 from NF-ΚB repression, and cell cycle re-entry, by interfering with the master regulation of caudal fin proliferation Fgf20a, thereby enabling OBs to act as progenitor cells. Moreover, glycolysis is necessary to maintain the correct proliferative ability and distribution of OBs populations within the blastema, during its formation. Glycolysis inhibition has a severe impact on OB dedifferentiation and pre-OBs pool assembly, which supports new OB formation and proliferation, ultimately leading to impaired bony-ray regeneration and suppression of blastema formation.

### Metabolic reprogramming in the caudal fin as a prelude to the stem cell state

Our results provide the first evidence that OB and other cell types respond to amputation by engaging metabolic routes that boost lactate-producing glycolysis instead of OXPHOS, thereby acquiring metabolic traits of stem cells. It is well described that embryonic stem cells and adult stem cells exhibit metabolic preferences distinct from their differentiated progeny. Both primed embryonic stem cells (26,35,36,86,87) and quiescent and proliferating adult stem cells(27,29,30,40,88) seem to rely on glycolysis, and once they differentiate they undergo a metabolic rewiring to increase mitochondrial biogenesis and OXPHOS. This reflects an essential role of glycolysis in periods of rapid cellular growth while oxidative metabolism is preferred in mature cells to maintain homeostasis(89, 90). Prioritizing glycolysis entails several advantages for rapid proliferating cells: fuels biosynthetic pathways necessary to sustain rapid cell growth and division by generating intermediaries for macromolecules synthesis (e. g., nucleic acids, lipids; and non-essential amino acids); the rate ATP generation is faster through glycolysis than mitochondrial glucose oxidation(39,42,43,59,91); and finally, pathways branching from glycolysis also provide intermediate metabolites necessary for post-translational modification of proteins, histones and DNA (e. g., acetylation, methylation, phosphorylation or glycosylation)(28,92–95). The latter, extends the connection between metabolism and modulation of intracellular signalling pathways, and the epigenome, to control gene expression programs that change cell function and fate(28,92–95). One of the best reported examples occurs during induced-pluripotent stem cell (iPSC) reprogramming, in which the switch towards a glycolytic metabolism happens before the expression of endogenous reprogramming factors(96, 97), implying that metabolic reprogramming is a cause rather that a consequence of cell reprogramming.

### Caudal fin metabolic reprogramming resembles the Warburg effect

Metabolic reprogramming has also emerged in a disease context as a cancer hallmark and first described in cancer cells as the “Warburg effect” (i. e., aerobic glycolysis) in which cancer cells use primarily glycolysis, resulting in lactate production, instead of pyruvate oxidation through OXPHOS(39–41,91). The metabolic switch that occurs during fin regeneration shares several parallels between cancer metabolism pathophysiology, namely preference for glycolysis to support proliferation and elevated levels of glutamine, an essential nutrient that supplies cancer metabolism. Besides functioning as a precursor for nucleotides and amino acid synthesis, glutamine can be converted into glutamate, a metabolic intermediate with various fates in proliferating cells (e. g., protein synthesis, and incorporation into the TCA)(98, 99). Interestingly, like cancer cells, our data points to an important role of glutamine and glutamate for the assembly of the blastema primordium, as our metabolome studies show an increase by 100- and 400-fold in glutamine and glutamate at the beginning of blastema formation, respectively. Cancer cells also produce high levels of lactate where it is often regarded as an important oncometabolite(100), correlated with cancer-induced angiogenesis, invasion, metastasis, and immunosuppression(101, 102). Our studies show not only an increase in lactate during the initial stages of regeneration, but also reveal that inhibition of pyruvate conversion to lactate leads to defects in blastema formation, although milder when compared to glycolysis inhibition. This indicates that lactate production may also contribute to proper blastema formation. The mechanisms by which the glutamine and glutamate cycle and lactate influence blastema formation should be addressed in future studies. Unexpectedly, albeit aerobic glycolysis is known to support cancer cell migration, our results show that glycolysis is not required for OB recruitment and motility. It would be interesting to evaluate how alterations in glucose metabolism are regulated throughout the regenerative process, without falling into tumorigenesis.

### Metabolic reprogramming is necessary for cell fate transitions and blastema proliferation

Our data shows that enhancing glycolysis serves to adapt to the cellular demands of regeneration, but it also seems to have the power to dictate several aspects of the regeneration program, induing modulation of mature OB dedifferentiation. Previous studies shown that OB dedifferentiation is a result of the activity of the NF-kB-RA axis(57, 67). In homeostasis, NF-kB supports RA signalling, by blocking the expression of *cyp26b1*, the RA-degrading enzyme, maintaining OB differentiation. After amputation, NF-kB becomes inactivated and Cyp26b1 suppression is lifted, thereby protecting OB from RA and promoting their dedifferentiation(57, 67). We show that blocking glycolysis leads to NF- kB signalling stimulation and decrease in *cyp26b1* expression, providing evidence that metabolic reprogramming precedes and is necessary to induce mature OB reprogramming into pre-OB. In addition, our data shows that glycolysis is necessary to support pre-OB cell cycle re-entry and to sustain blastema proliferation (Fig 7) and can be linked to *fibroblast growth factor 20a* (*fgf20a*), which is fundamental for blastema initiation and proliferation during regeneration(72,75,103). Mutants for *fgf20a* fail to form a functional blastema and are unable to proliferate(103). Accordingly, blocking Fgf receptor 1 activity leads to a similar phenotype(71, 72), but does not impair OB dedifferentiation(8), suggesting that its primary role is to regulate blastema proliferation. Our work suggests that glycolysis promotes not only the expression of *fgf20a*, but also of other ligands that cooperate to induce *fgf20a* expression, such as *igf2b* and *wnt10a*(73, 74). Since these pathways are part of a general mechanism triggered upon amputation to stimulate proliferation, it is not surprising that glycolysis inhibition caused an overall reduction of proliferation. In addition, the presented data indicates that glycolysis is necessary until the end of blastema formation, to generate new OBs and to maintain a proper balance between OB subtypes within the blastema (Fig 7). Decline in the total number of immature OB and in the proliferative rates of distal pre-OB and of proximal immature OB populations observed upon glycolysis inhibition, can be accounted, at least in part, by the pronounced effects of glycolysis inhibition on blastema proliferation. Importantly, these results corroborate the idea that regeneration benefits from glycolysis both in terms of biomass generation, to support cell proliferation, but also by inducing the expression of powerful mitogens, like *fgf20a*. Noteworthy, besides mature OBs, pre-OB can also derive from a population of OB progenitor that resides in the joint regions of the fin(58). Thus, mature OB and joint-associated progenitors may act as complementary sources that supply the pre-OB pool. In fact, we observed that glycolysis inhibition leads to a diminished number of pre-OBs before blastema formation, yet this number is back to normal after blastema formation. Since blocking glycolysis prevented OB dedifferentiation, we could speculate that over time OB progenitors from the joints where able to replenish the pre-OB pool. Further work is needed to test this hypothesis and the impact of metabolic reprogramming in supporting OP activation and contribution for new OB formation. In general terms, this study provides the first line of evidence that metabolic reprogramming towards glycolysis governs mature OB plasticity and blastema proliferation. To some extent, this is mediated through glycolysis-driven changes in gene expression that allows to uncouple dedifferentiation from acquisition of proliferative capacity.

### Metabolic reprogramming as a conserved mechanism in regenerative contexts

Regeneration is in its essence an anabolic process. After an insult, reconfiguration of the extracellular milieu can induce metabolic adaptations that are fundamental to accommodate new cellular functions that support growth and cell fate decisions necessary for regeneration. In line with our data, other animals with enhanced regenerative abilities, such as planarians(48) or amphibians(50,51,104) also show a predominance of glycolysis upon injury to sustain proliferation. This indicates that metabolic rewiring towards glycolysis might be a conserved mechanism necessary for the regenerative process. Importantly, metabolic reprogramming was also shown to be necessary for regeneration of other zebrafish tissues. Like OBs in the fin, after cardiac injury, regeneration is achieved via dedifferentiation and proliferation of cardiomyocytes near the injury(105, 106). Recent studies have demonstrated that these cells switch to a glycolytic metabolism necessary for their dedifferentiation and proliferation(53, 54). Moreover, regeneration of the embryonic tail was shown to rely on glycolysis to support blastema formation by (52). Glycolysis was required to fuel the hexosamine pathway(52), which is responsible for glycosylation of proteins associated with cell signalling, gene transcription and EMT(107, 108). Given these results and the similarities between larval tail and adult caudal fin regeneration, it would be important to examine the function of hexosamine pathway during fin regeneration. In contrast to zebrafish, mammals possess poor capacity to perform epimorphic regeneration of complex structures, with only a few examples, such as amputated ear and digit tips(24, 25). In mice models of ear and digit injuries, regeneration is impaired by OXPHOS inhibition, suggesting that in this context OXPHOS is required to mediate regeneration(109). In contrast, the MRL mice strain, which has an enhanced regenerative capacity in comparison to other mice, showed an increase of aerobic glycolysis over OXPHOS after injury of several organs(110, 111). This indicates that further studies are necessary to clarify the potential role of glucose metabolism during mammalian regeneration. In regard to bone, disruption of the metabolic profile of OBs and OB sources (e. g., mesenchymal stem cells) might also have important implications for bone repair after injury and in certain pathological conditions (e. g., osteoporosis), as they influence OB identity status and function(44,100,112,113). Cell metabolism can potentially be a target in the contexts of fracture healing or bone diseases, to stimulate the repair process, or to prevent OB dysfunction.

## Concluding remarks

The data described here provides the first evidence that a metabolic reprogramming favouring glycolysis over mitochondrial oxidation occurs at early stages of adult regeneration and is an integral component of the regenerative program. This is in accordance with recent regeneration studies performed in other systems and resembles many traits of the Warburg effect observed in cancer cells. Our data indicates that OB and possibly other cell lineages, favour aerobic glycolysis, to engage a specialized genetic program that enables them to act as progenitor cells. We unveil a novel and fundamental role of glycolysis in mediating mature OB dedifferentiation and cell cycle re-entry and supporting blastema assembly and proliferation. Moreover, we have uncoupled the effects of glycolysis in mediating OB dedifferentiation from proliferation by identifying distinct downstream transcriptional targets of the glycolytic metabolism. This provides evidence that the role of metabolic reprogramming in regeneration is not limited to sustain macromolecule synthesis and energy production. Overall, our findings support a notion that glucose metabolism has a powerful instructive role in regulating lineage-specific programs and generic responses to injury that induce changes in cell identity and function, crucial to prompt bone regeneration.

## Material and Methods

### Ethics statement

All the people involved in animal handling and experimentation were properly trained and accredited by FELASA. All experimental procedures were approved by the Animal User and Ethical Committees at Centro de Estudos de Doenças Crónicas (CEDOC) and accredited by the Portuguese National Authority for Animal Health (DGAV), according to the directives from the European Union (2010/63/UE) and Portuguese legislation (Decreto-Lei 113/2013) for animal experimentation and welfare.

### Zebrafish lines maintenance and caudal fin amputation

Wild-type AB and transgenic zebrafish lines, namely *Tg(osterix:mCherryNTRo)^pd4^* (114) (referred as *osx*:mCherry), kindly provided by Kenneth Poss, *Tg(ola.Bglap:EGFP)^hu4008^* (referred as *bglap*:EGFP) and *Tg(Has.RUNX2-Mmu.Fos:EGFP)^zf259^* (8) (referred as *runx2*:EGFP), kindly provided by Gilbert Weidinger, were maintained in a circulating system with 14-hour/day and 10-hour/night cycle at 28°C(115). All regeneration experiments were performed in 4-18 months-old fish and transgenics used as heterozygotes. Caudal fin amputations were performed in fish anaesthetized with buffered 160 mg/mL MS-222 (Sigma, E10521), using a sterile scalpel to remove approximately one half of the fin, as previously described(72). Fish were left to regenerate in an incubator at 33°C ± 1°C with water from the circulating system until defined time-points. Regenerated fins were collected from anaesthetized fish, and either processed for cryosectioning, stored in Trizol for RNA isolation, handled for Mass- spectrometry (MS) or for flow cytometry.

### Pharmacological and chemical treatments

For pharmacological treatments via intraperitoneal injections (IP), fish were randomized and subjected to IP injections at the designated time-points, with either 2DG (Sigma-Aldrich, 0,5 mg/g diluted in 1x Phosphate Buffered Saline (PBS)), S.O. (Sigma-Aldrich, 0,6 mg/g diluted in 1x PBS), UK- 5099 (Sigma-Aldrich, 0,02 mg/g, diluted in a mixture of 1x PBS and DMSO (1:1)) or with corresponding vehicle (Control). IP injections were performed with an insulin syringe U-100 G 0,3 mL and a 30G needle (BD Micro-fine) inserted close to the pelvic girdle. For 3PO (Sigma-Aldrich) and MB-6 (Calbiochem) treatments, compounds were diluted in DMSO and added to water from the circulating system to a final concentration of 15µM and 2,5 µM, respectively, and controls with equivalent amount of vehicle. For all experiments water was replaced daily and fish left to regenerate until the desired time-point.

For glucose uptake assay, fish were administered with the glucose analogue 2-NBDG (Sigma-Aldrich, 25 µmol/kg, from stock solution dissolved in DMSO) via IP injection 1h prior to imaging. For S-phase labelling, fish were subjected to caudal fin amputation and administrated with Ethynyl-2’-deoxyuridine (EdU, Thermo Scientific: C10337, 20 µL of 10 mM solution diluted in 1x PBS) via IP injection 3h prior to caudal fin collection.

### Total RNA isolation and quantitative real-time PCR (qPCR)

For gene expression analysis, caudal fin composed of the regenerated tissue and one bony-ray segment proximal to the amputation plane were collected. Pools from 4-5 caudal fins were used per biological replicate. Briefly, samples were homogenized in Trizol reagent (Invitrogen, 15596026) for cell disruption and RNA extracted as previously described(13), using the RNeasy Micro kit (Qiagen, 74004) according to manufacturer’s protocol. cDNA was synthesized from 1 μg total RNA for each sample using the Transcriptor High Fidelity cDNA Synthesis Kit (Roche, 05081963001), with a mixture of oligo dT and random primers. All qPCR primers are listed in Sup Table 1. qPCR was performed using a FastStart Essential DNA Green Master Mix (Roche, 4385617) and a Roche LightCycler 480. Cycle conditions were: 15 min pre-incubation at 95°C and 3 step amplification cycles (45x), each cycle for 30 sec at 95°C, 15 sec at 60°C and for 30 sec at 72°C.

### Alizarin Red S staining and immunofluorescence in cryosections

*In vivo* Alizarin red S (ARS, Sigma-Aldrich) staining in the *bglap*:EGFP transgenic was performed prior to caudal fin amputation as previously described(116). Briefly, fish were incubated in a 0.01 % ARS solution, dissolved in water from the circulating system and pH adjusted to 7.4 with a KOH solution, for 15 min in the dark and rinsed 3 times, 5 min each. Caudal fins were amputated and imaged at specific time-points post-amputation.

Tissue processing for cryosections was performed as previously described(13). Shortly, fins were collected, fixed overnight (ON) in 4% paraformaldehyde (in 1x PBS) and stored in 100% methanol (MeOH) at -20°C, until subsequent analysis. They were then gradually rehydrated in a series of MeOH/1x PBS (75%, 50% and 25%) and incubated ON in 30% sucrose (Sigma-Aldrich, diluted in 1x PBS). For EdU labelling, caudal fins were fixed and directly incubated in 30% sucrose solution. Fins were then embedded in 7.5% gelatin (Sigma-Aldrich)/ 15% sucrose in 1x PBS and subsequently frozen in isopentane at -70°C and stored at -80°C. Longitudinal caudal fins sections were obtained at 12 μm using a Microm cryostat (Cryostat Leica CM3050 S) and slides stored at -20°C. For immunofluorescence on cryosections, slides were thawed for 15 min at room temperature (RT), washed twice in 1x PBS at 37°C for 10 min and subjected to an antigen retrieval step, which consisted of a 15 min incubation at 95°C with sodium citrate buffer (10mM Tri-sodium citrate with 0.05% Tween20, pH 6). Slides were then incubated in 0.1 M glycine (Sigma-Aldrich, in 1x PBS) for 10 min, permeabilized in acetone for 7 min at -20°C and incubated for 20 min in 0.2% PBST (1x PBS with 0.2% Triton X-100). At this point, cryosections used for EdU labelling were incubated with the labelling solution according to the manufacturer’s protocol (Thermo Scientific: C10337). For TUNEL labelling assay, cryosections were permeabilized in a sodium citrate solution (0.1% sodium citrate and 0.1% Triton X-100 in 1x PBS) and labelled according to the manufacturer’s protocol (Roche, 11684795910). Afterwards, they were incubated in a blocking solution of 10% non-fat dry milk in PBST for 2-4 h at RT. Slides were then incubated with primary antibodies diluted in blocking solution, ON at 4°C (for antibody details see Sup Table 2). On the following day, slides were washed with PBST 6 times, 10 minutes each, and incubated with secondary antibodies (Sup Table 3) diluted in blocking solution, for 2 h at RT and protected from light. Subsequently, slides were washed 3 times, 10 min each, in PBST and then counterstained with 4’,6-diamidino-2-phenylindole (DAPI; 0.001 mg/mL in 1x PBS, Sigma-Aldrich) for 5 min in the dark, for nuclei staining. Slides were then washed 3 times with PBST, 10 min each, mounted with fluorescent Mounting Medium (DAKO) and stored at 4°C protected from light until image acquisition.

### Flow cytometry

For fluorescence-activated cell sorting (FACS) of OB, caudal fins from *bglap*:EGFP transgenic line were amputated, tissue collected at specific time-points during regeneration and dissociated into single cell suspensions. For that, fins were incubated for 20 min at 28°C with vigorous shaking in a solution of Liberase DH Research Grade (0,05mg/ml in 1x PBS, Roche). Cell suspensions were passed through a 30 μm filter (CellTricks, Sysmex) and centrifuged at 300g for 5 min at 4 °C. Cell pellets were resuspended in 1x PBS with 10% fetal bovine serum (Biowest). FACS was carried out on a MoFlo high-speed cell sorter (Beckman Coulter, Fort Collins, USA) using a 488 nm laser (200 mW air-cooled Sapphire, Coherent) at 140 mW for scatter and a 530/40 nm bandpass filter for GFP measurements. Cell debris and aggregates were removed from the analysis. The fluorescence scatter (Comp-FL Log::GFP) was used to separate cells according to their GFP fluorescence intensity with a maximum of stringency to avoid cross-contamination. Zebrafish WT AB strain was used as a negative control to set the GFP-positive population. The instrument was run at a constant pressure of 207 kPa (30 psi) with a 100 µm nozzle and frequency of drop formation of approximately 40 kHz. Two and three independent biological replicates were performed for each condition at 0 and 6 hpa respectively. For each, 300 GFP-positive cells were collected directly into lysis and RNA stabilization buffer (provided by OakLabs GmbH) and vigorously shaken for 1 min. To verify the quality of the samples, cell death and purity were measured. Cell death was measured by incubating the samples with propidium iodide (PI, Sigma-Aldrich), to a concentration of 1 μg/ml, and using the 488 nm laser for PI excitation and measure on the PI channel (613/20 BP). Only samples with cell death below 10-20% and purity above 90% were used for subsequent analysis. Samples were maintained at -80°C until sent to OakLabs GmbH (Henningsdorf, Germany) for cDNA generation, microarray chip set up and data analysis.

### Osteoblast ArrayXS

To compare the transcriptome profiles of mature OB in homeostasis to OB during dedifferentiation, a genome-wide gene expression profiling was set up using the 8×60K ArrayXS Zebrafish platform by Agilent and performed by OakLabs GmbH (Henningsdorf, Germany). The 8×60K ArrayXS Zebrafish represents approximately a total of around 60000 zebrafish transcripts, which includes 48000 coding genes, 8075 non-coding genes and 19140 predicted genes annotated in the Zv9 release 75. RNA quality was processed by Oaklabs using the 2100 Bioanalyzer (Agilent Technologies), the RNA 6000 Pico Kit and a photometrical measurement with the Nanodrop 2000 spectrophotometer (Thermo Scientific). Sample quality was evaluated based on the Bioanalyzer’s RNA integrity number (RIN). Only samples with RIN ≥ 8 were used. Subsequently, 2 µL of the lysis and RNA stabilization buffer, from three biological replicates of each condition (0 and 6 hpa OBs), was used for cDNA synthesis and pre-amplification using the Ovation One Direct system (NuGEN). The generated cDNA was labelled with Cy3--dCTP using the SureTag DNA Labelling Kit (Agilent) prior to microarray hybridisation. Microarray blocking, hybridisation and wash were performed using Agilent’s Oligo aCGH/ChIP-on-Chip Hybridisation Kit, following the manufacturer’s protocol. Ultimately, fluorescence signals were detected by the SureScan Microarray Scanner (Agilent Technologies), at a resolution of 3 µm for SurePrint G3 Gene Expression Microarrays and 5 µm for HD Microarray formats. This resulted in a raw data output of 1-colour hybridisation using the Agilent’s Feature Extraction software version 11. Raw data was then subjected to processing and analysis. Briefly, background signals were subtracted and then normalized using the ranked mean quantiles(117). For data quality control and to identify potential outlier samples, hierarchical clustering and a principal component analysis were performed. The retrieved data was used to compare the expression profiles of OB from 6 hpa with 0 hpa. Accession number for the transcriptome data sets is GSE194385.

### Liquid chromatography-mass spectrometry (LC-MS) analysis

For metabolite analysis, caudal fins were collected and snap-freeze in liquid nitrogen for 5 minutes and diluted in a mixture containing MeOH:dH2O (2:1) and an internal standard α-Aminobutyric acid (AABA, 2mM final concentration). Samples were homogenized using tissue grinder for 5 seconds and using the ultrasound bath for 30 minutes at 4°. This was followed by sample centrifugation for 10 minutes at top speed at 4°, supernatant collected and stored at -20° (short storage) or -80°C (long storage). Samples and internal standards were analysed in a Dionex UltiMate 3000 UHPLC (Ultra-High Performance Liquid Chromatography) system coupled to a heated electrospray QExactive Focus mass spectrometer (Thermo Fisher Scientific, MA, USA). Three separate LC-MS assays were applied. For glutamine (Duchefa) and glutamate (Thermo Fisher Scientific) detection, we used an Acquity UPLC BEH Amide column and spectra were acquired in positive ionization mode using a method which consisted of several cycles of FullMS scans (75-1125 m/z; Resolution=70 000 FWHM at 200 m/z). Glucose (Sigma Aldrich) and lactate (Alfa Aesar) detection were performed with acquisition in negative ionization mode. For every assay, 4 biological replicates (10 fins used per replicate) were used per condition and sample injection was performed in triplicate and a volume of 5 µL was applied.

### Image acquisition and processing

For regenerated area measurements, images of live anesthetised WT and transgenic adult caudal fins were acquired in a Zeiss Lumar V-12 fluorescence stereoscope equipped with a Zeiss axiocam MRc camera using a 0.8X air objective (at 14x zoom) controlled by Zen 2 PRO blue software. Images were acquired using transmitted light and the GFP (FS05) and/or TexasRed (FS45) filters, according to the fluorescent reporter expressed. Images were assembled using the Fiji software(118).

For 2-NBDG labelled WT caudal fins, images were acquired using in a Zeiss Axio Observer z1 inverted microscope for transmitted light and epifluorescence, equipped with an axiocam 506 monochromatic camera, using an EC Plan-Neofluar 5x 0.16NA air objective controlled by Zen 3 blue software. An image mosaic was acquired using transmitted light and the GFP (38HE) filter. Serial sections were acquired every 5 µms. For image processing, composite maximum intensity images and concatenation of several images along the caudal fin proximal-distal axis was performed using the Zen 3 blue software and images assembled using Fiji software(118).

For live-imaging analysis of OB migratory dynamics *in vivo*, *bglap*:EGFP transgenic fish were anesthetised and maintained in glass bottom Petri dishes. Imaging was performed in a confocal microscope Zeiss LSM 710 using the software ZEN 2010B SP1. Fish were imaged with a Plan-Neofluar 10x 0.3 NA air objective using the 488 nm (emission windows:490-530nm) and 568 nm (emission windows:570-650nm), if counterstained with ARS, excitation wavelengths coupled with a transmitted light PMT. Serial sections were acquired every 5 µms. For OB motility assay, time-lapse images were acquired always in the same region of the fin, capturing the first 2 segments below the amputation plane (segment 0 and segment -1) and the blastema region, and images acquired every 5 h following amputation, during the first 25 hpa. For assessment of OB migration in vehicle and 2DG treated fish, time-lapse images were acquired at 0 and 24 hpa. For image processing, composite maximum intensity z-stack projections were made using the Fiji software(118). Time-lapses were assembled and computationally registered with the Fiji StackReg and MultiStackReg plugins(118).

Immuno-labelled cryosections were analysed in confocal microscopes Zeiss LSM 710 and Zeiss LSM 980 controlled by ZEN 2010B SP1 or ZEN 3.3, respectively. Cryosection images were acquired using a C-Apochromat 40x 1.2NA water objective with 0.6x zoom, a step size of 1 µm, and 405, 488, 568, and 633 nm excitation wavelengths coupled with transmitted light PMT. Sequential images were acquired to capture the first segment below the amputation plane and the entire regenerated region. For image analysis and processing, composite maximum intensity z-stack projections were made using the Fiji software(118). When required, concatenation of several images along the proximal-distal axis of the same longitudinal section was performed using the Fiji plugin 3D Pairwise Stitching(118).

For all cryosections of manipulated fish and corresponding control and time-lapse assays, images were acquired employing identical settings (magnification, contrast, gain and exposure time) and in identical/comparable regions. All Images were then processed using the Adobe Photoshop CS5 and Adobe Illustrator CC.

### Quantifications and statistical analysis

For qPCR analysis, all samples were analysed in four to eight biological pools. For each biological pool, qPCR was performed for each target gene in 3 technical replicates. Gene expression values were normalized using the *elongation factor 1α* (*ef1α*, NM_131263) housekeeping gene and relative expression was calculated using the 2(-ΔΔC(T) method(119).

Measurements of total regenerated area were obtained by delineating the fin regenerated area using the Area tool in Fiji. The regenerated area was then normalized to the corresponding total caudal fin width to avoid discrepancies related to the animal size, resulting in one measurement per animal.

Total number of pre-OB (Runx2^+^) and OB subtypes (Runx2^+^Bglap^+^ or Runx2^+^Osx^+^) in longitudinal cryosections was quantified by analysing the number of labelled cells in relation to the imaged cryosection area (per 100 µm^2^), determined using the Area tool on Fiji. Percentage of pre-OB and OB subsets was quantified by analysing the number of cells in the imaged cryosection area in relation to the total number of OB lineage cells (Runx2^+^ and Runx2^+^Bglap^+^or Runx2^+^ and Runx2^+^Osx^+^).

Percentage of proliferating Bglap^+^ OBs was quantified by analysing the number of Bglap^+^ PCNA^+^OBs in the imaged cryosection area in relation to the total number of Bglap^+^ OBs, per time-point analysed. Total number of EdU^+^ pre-OB (Runx2^+^) and OB subtypes (Runx2^+^Bglap^+^ or Runx2^+^Osx^+^) was quantified by analysing the number of labelled cells in relation to the imaged cryosection area (per 100 µm^2^). Percentage of EdU^+^ pre-OB (Runx2^+^) and OB subtypes (Runx2^+^Bglap^+^ or Runx2^+^Osx^+^) was quantified by analysing the number of EdU^+^ cells within the cryosection area in relation to the total number of OB lineage cells (Runx2^+^ and Runx2^+^Bglap^+^or Runx2^+^ and Runx2^+^Osx^+^).

Total number of TUNEL^+^ and EdU^+^ cells was assessed by quantifying the number of labelled cells within each fin compartment (mesenchyme and epidermis), in relation to the corresponding compartment area (per 100 µm^2^), determined using the Area tool on Fiji.

All quantifications were done using the Cell-counter plugin on Fiji in individual cryosections representing at least 3 different blastemas per animal and 3-5 animals per condition.

For quantification of OB motility during regeneration, live-imaging time-lapses of bony-rays, including the segment 0 and segment -1, were used. Quantification was performed using a custom Matlab script that performs all the workflow. Both GFP and brightfield (BF) channels were gaussian filtered (sigma 2) to reduce noise. Intersegment regions were found to define the boundaries between the segments analysed using the BF channel and a sobel vertical algorithm, dilated with a vertical kernel and small connected components (200pixels) removed resulting in an average line profile for each bony-ray. Intersegments (peaks) were found using findpeaks matlab function. OB location was tracked by finding the global GFP intensity center (center of mass) in segment 0 and -1. GFP line profiles were calculated and summed in height and the intensity center of mass was found in each segment analysed. The final result is expressed as a ratio (relative OB displacement) between the center of mass location and the total segment length (0: Anterior bias; 1: Posterior bias).

For LC-MS, raw data was analysed using Xcalibur’s Quan Browser (version 4.1.31.9, Thermo Scientific). The peaks corresponding to each compound of interest were identified by comparison with standards analysed in the same conditions. A mass tolerance of 5 ppm and a retention time window tolerance of 10 sec were used. Peak areas used for relative quantitation were obtained using the Genesis method. Peak detection considered the nearest peak to the retention time defined for each compound, with a minimum peak height (S/N) of 10. The peak area from AABA was used as an internal quantitation calibrant for the final quantitative data. Quantitation variability was assessed by the calculation of the relative coefficient of variance (CV %).

Statistical analysis was performed in GraphPad Prism v7 and statistical significance considered for *p* < 0.05. Statistical tests, P values, mean and error bars are indicated in the respective figure legends. For sample size see Sup table 4. For OB ArrayXS, fold change was determined based on the normalised data set and expression ratios obtained. In this data set, a logarithmic base 2 transformation was performed (i.e. log2 (expression ratio)) to make the mapping space symmetric and the up-regulation and down-regulation comparable, prior to the significance test. The mean and standard deviations of the two sets of isolated OB samples (0 hpa and 6 hpa) were then compared using a Welch’s t-test (or unequal variances t-test), generating 1 data set with the differential expressed genes between both conditions. Only p-values less than 0,05 were considered statistically significant and log2 values that lie between -1 and 1 were ignored.

## Acknowledgements

We are grateful to Lara Carvalho for reading the manuscript. Ana Teresa Tavares and Carolina Crespo for useful discussion and the Tissue Repair and inflammation Lab for support. We acknowledge Ana Farinho from CEDOC’s Histology Facility for assistance in tissue processing and cryosectioning. We thank Petra Pintado and Fábio Valério from CEDOC’s Fish Facility for technical assistance with the support from the research consortia Congento, co-financed by Lisboa Regional Operational Program (Lisboa2020), under the Portugal 2020 Partnership Agreement, LISBOA-01-0145-FEDER-022170. We thank the technical support from CEDOC’s Microscopy Facility, funded by PPBI-POCI-01-0145-FEDER- 022122. To the Instituto Gulbenkian de Ciência’s Flow Cytometry facility and Cláudia Andrade for technical support. ASB, JB, RL and AJ thank Fundacão para a Ciência e a Tecnologia for funding, PTDC/BIA-BID/29709/2017 and PTDC/BTM-SAL/29377/2017. ABB thanks ATIP-Avenir 2020 for funding. RL is supported by Fundação para a Ciência e Tecnologia, in the context of a program contract to RL (4, 5, and 6 of article 23° of D.L. no. 57/2016 of August 29, as amended by Law no. 57/2017 of 19 July).

---

**Sup Fig 1.**
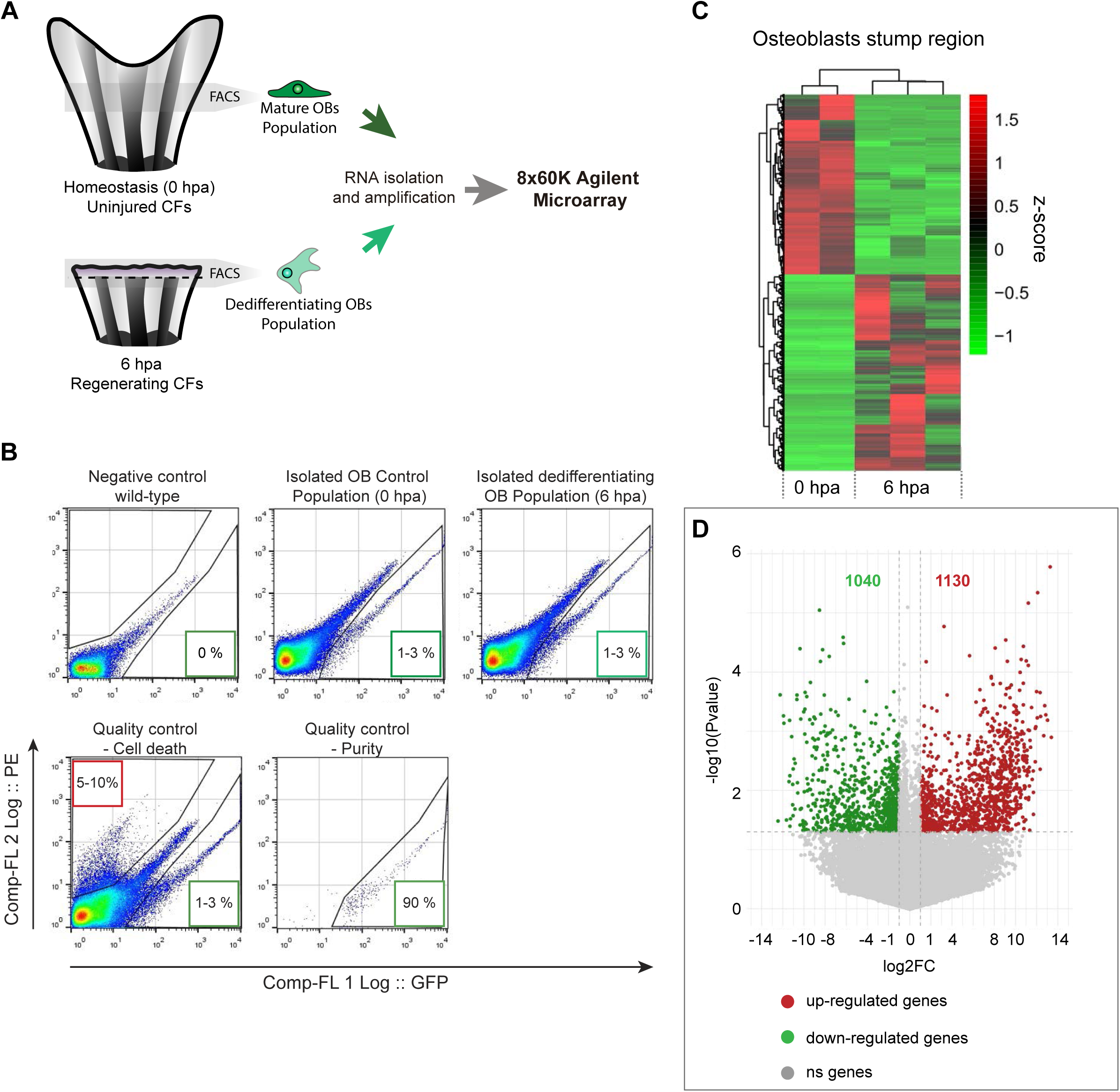
Isolation and gene expression analysis osteoblasts undergoing dedifferentiation. (A) Schematic representation of the experimental design used to obtain the transcriptional profile of OBs undergoing dedifferentiation, using bglap:EGFP reporter line. OBs from caudal fin tissue corresponding approximately to one bony-ray segment (grey areas), from homeostasis (0 hpa, corresponding to mature OBs) and from one segment bellow amputation at 6 hpa (dedifferentiating OBs) were collected, dissociated, and isolated by FACS for RNA extraction and subjected to a microarray chip assay. (B) Representative flow cytometry plots from wild-type (negative control) and bglap:EGFP transgenic fish from homeostasis and 6 hpa caudal fins. Representative examples of sample quality control through the evaluation of cell death, by propidium iodide (PI) staining, and purity. In flow cytometry plots, GFP fluorescence intensity is given by the x axis (Comp-FL 1 Log ::GFP) and PE fluorescence intensity (used to identify PI-positive cells) is given by the y axis (Comp-FL2 Log::PE). Numbers in the lower right boxes indicate relative percentages of GFP+ cells and numbers in the upper left boxes indicate the relative percentage of PI+ cells. (C) Heatmap depicting hierarchical clustering of Z score transformed expression profiles of all genes from isolated osteoblasts at 0 and 6 hpa. (D) Volcano plot showing the differentially expressed transcripts between homeostatic (6 hpa) and dedifferentiating osteoblasts (0 hpa). The horizontal axis represents the log2 fold-change between 6 and 0 hpa and the vertical axis represents the p-value (on a negative log10 scale). Upregulated transcripts are shown in red and downregulated genes are shown in green. Significant changes were considered for a log2 FC>1 or <-1 for a p-value> 0.05.

**Sup Fig 2.**
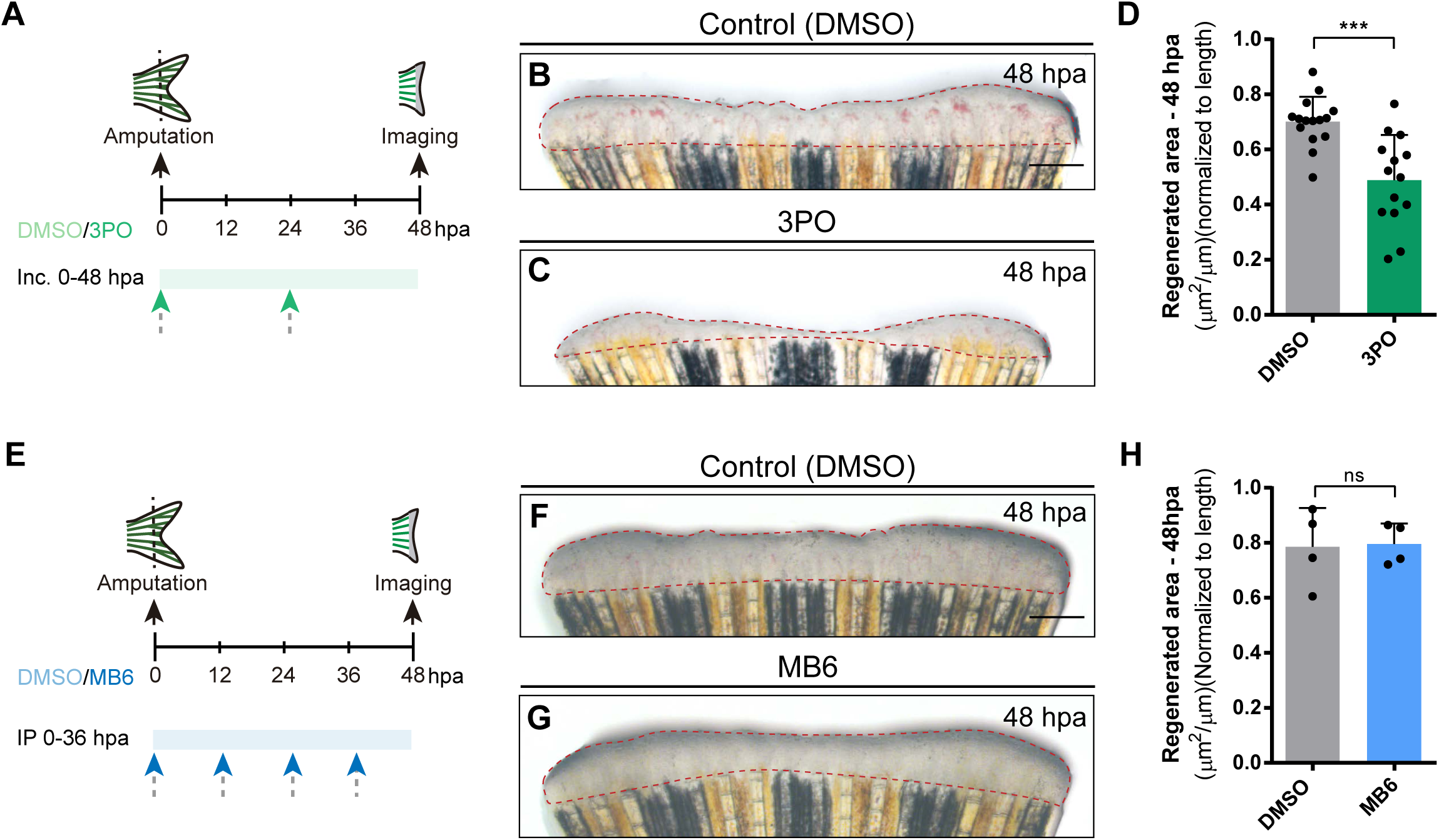
Inhibition of glycolysis impairs blastema formation. (A) Experimental design used to inhibit glycolysis during fin regeneration. Fish are incubated with control vehicle (PBS) or with the glycolytic inhibitor 3PO, every 24 hours from caudal fin amputation (0 hpa) until 48 hpa. (B-C) Representative images of 48 hpa fins treated with (B) vehicle (DMSO) or (C) 3PO. (D) Quantification of the total caudal fin regenerated area at 48 hpa of fish treated with vehicle (DMSO) or with 3PO. (E) Experimental design used to inhibit pyruvate mitochondrial import during fin regeneration. Fish are incubated with vehicle (DMSO) or MB6, every 24 hours from fin amputation (0 hpa) until 48 hpa. (F-G) Representative images of 48 hpa caudal fins treated with (F) vehicle (DMSO) or (G) MB6. (H) Quantification of the total caudal fin regenerated area at 48 hpa of fish treated with vehicle (DMSO) or with MB6. Statistical analysis displayed on the graphs corresponds to Mann-Whitney test with Mean ± SD, each dot corresponds to one fish. Scale bar represents 500 µm. Dashed lines define the regenerated area. ns: not significative; ***P<0,001.

**Sup Fig 3.**
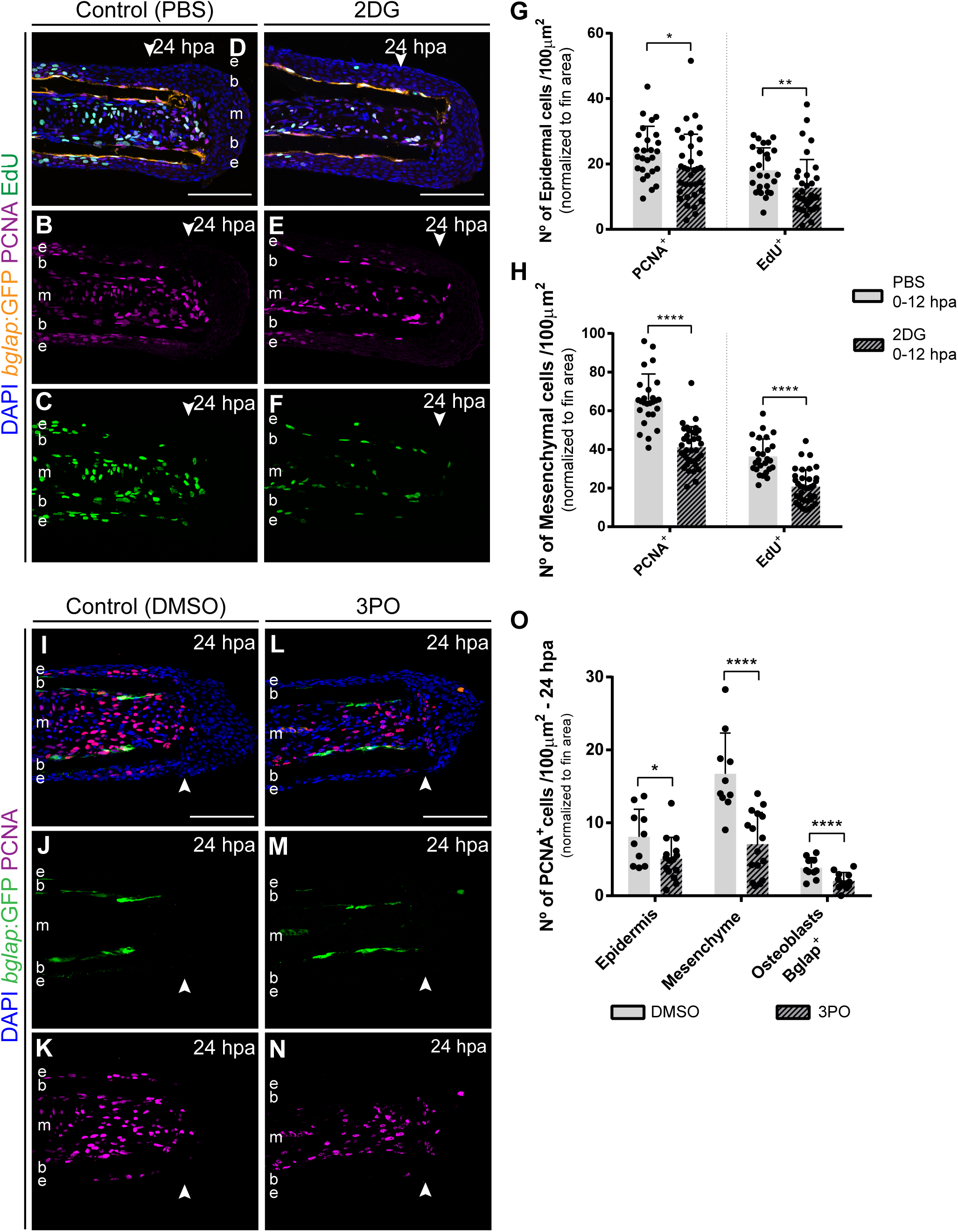
Inhibition of glycolysis prevents cell cycle re-entry. (A-F) Representative 24 hpa cryosection images of bglap:EGFP (orange) caudal fins immunostained for PCNA (magenta), EdU (green) and counterstained with DAPI (blue), in fish treated with (A-C) vehicle (PBS) or (D-F) 2DG. (G-H) Quantification of the number of PCNA+ proliferating cells and EdU+ proliferating cells at 24 hpa, in the (G) epidermis and (H) mesenchyme, in caudal fins from control and 2DG treated fish. Statistical analysis displayed on the graphs corresponds to Mann-Whitney test (n=26 (PBS) and 35 (2DG) cryosections). (I-N) Representative 24 hpa cryosection images of bglap:EGFP (green) caudal fins immunostained for PCNA (magenta) and counterstained with DAPI (blue), in fish treated with (I-K) vehicle (DMSO) or with (L-N) 3PO. (O) Quantification of the number of PCNA+ proliferating cells in the epidermal and mesenchymal compartments, and PCNA+ Bglap+ cells, at 24 hpa in fins from control and 3PO treated fish. Statistical analysis displayed on the graphs corresponds to Mann-Whitney test with Mean ± SD (n=10 (DMSO) and 14 (3PO) cryosections). Arrowheads indicate amputation plane. Scale bars represents 100 µm. e: epidermis; b: bone; m: mesenchyme. *P <0.05; **P<0,01; ****P<0,0001.

**Sup Fig 4.**
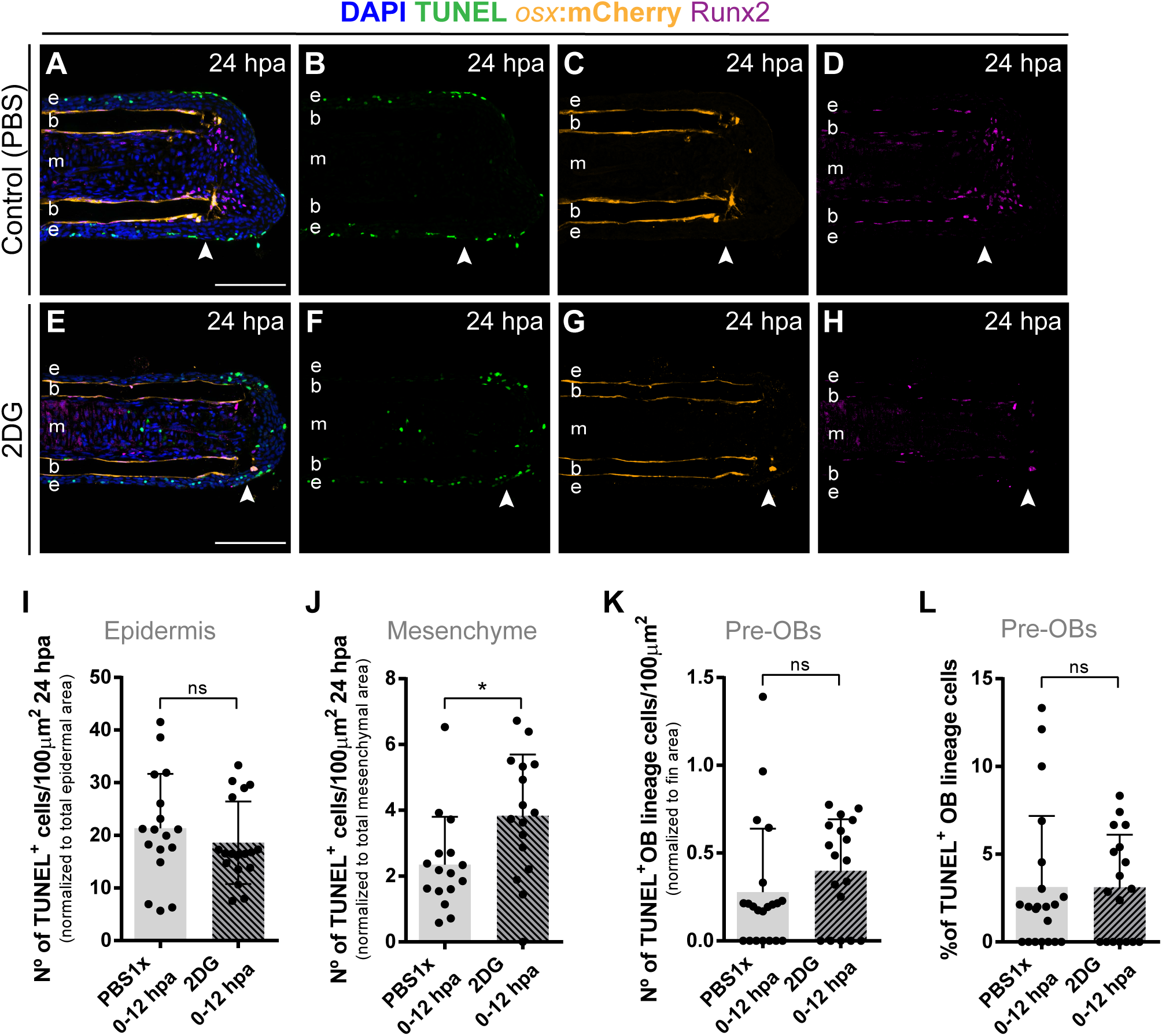
Inhibition of glycolysis has no effect on pre-osteoblasts cell death. (A-H) Representative 24 hpa cryosection images of osx:mCherry (orange) caudal fins immunostained for Runx2 (magenta), TUNEL (green) and counterstained with DAPI (blue), in fish treated with (A-D) vehicle (PBS) or (E-H) 2DG. Arrowheads indicate amputation plane. Scale bar represents 100 µm. e, epidermis; b, bone; m, mesenchyme. (I-K) Quantification of the total number of TUNEL+ cells in the (I) epidermis, (J) mesenchyme and (K) and in the pre-OBs at 24 hpa, in caudal fins from controls and 2DG treated fish. (L) Percentage of TUNEL+ pre-OBs at 24 hpa, in fins from controls and 2DG treated fish. Statistical analysis displayed on the graphs corresponds to Mann-Whitney test with Mean ± SD (n=16-21 cryosections). ns: not significant; *P <0,05.

**Sup Fig 5.**
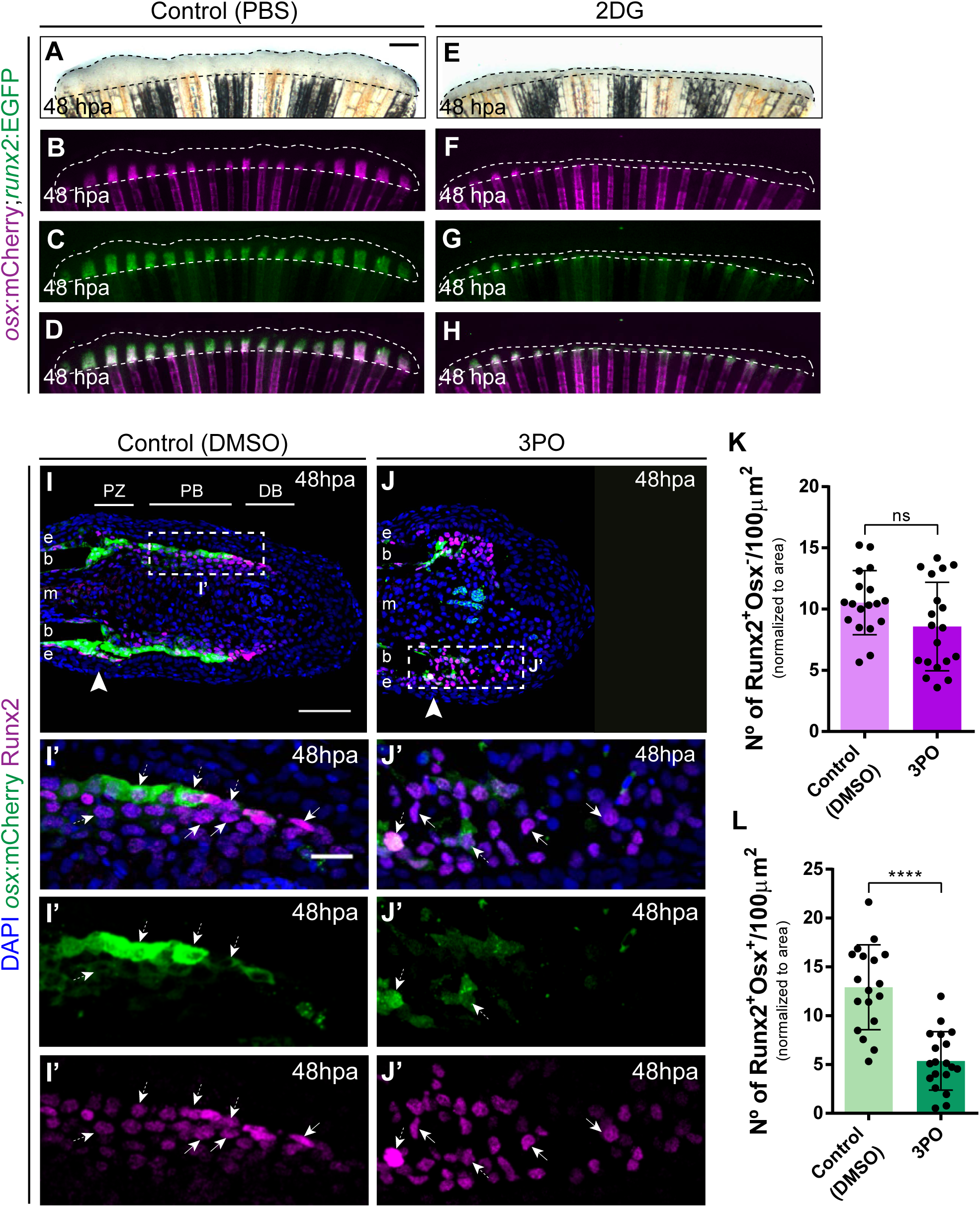
Inhibition of glycolysis affects distribution of osteoblast subtypes in the blastema. (A-H) Representative 48 hpa caudal fin images of osx:mCherry (magenta) and runx2:EGFP (green) double transgenic treated with (A-D) vehicle (PBS) or (E-H) 2DG. Dashed lines delineate regenerated area. Scale bar represents 500 µm. (I-J’) Representative 48 hpa cryosection images of osx:mCherry (green) caudal fins immunostained for Runx2 (magenta) and counterstained with DAPI (blue), in fish treated with (I,I’) vehicle (DMSO) or (J,J’) 3PO. Dashed boxes represent magnified panels in I’ and J’. Arrowheads indicate amputation plane. Scale bar represents 100 µm and 20 µm for magnified panels. PZ: patterning zone; PB: proximal blastema; DB: distal blastema. (K-L) Quantification of the number of (K) Runx2+Osx- and (L) Run2+Osx+ OB subtypes at 48hpa, in caudal fins from control and 3PO treated fish. Statistical analysis displayed on the graphs corresponds to Mann-Whitney test with Mean ± SD (n=18 (DMSO) and 19 (3PO) cryosections) ns: not significant; ****P <0,0001.

**Sup Fig 6.**
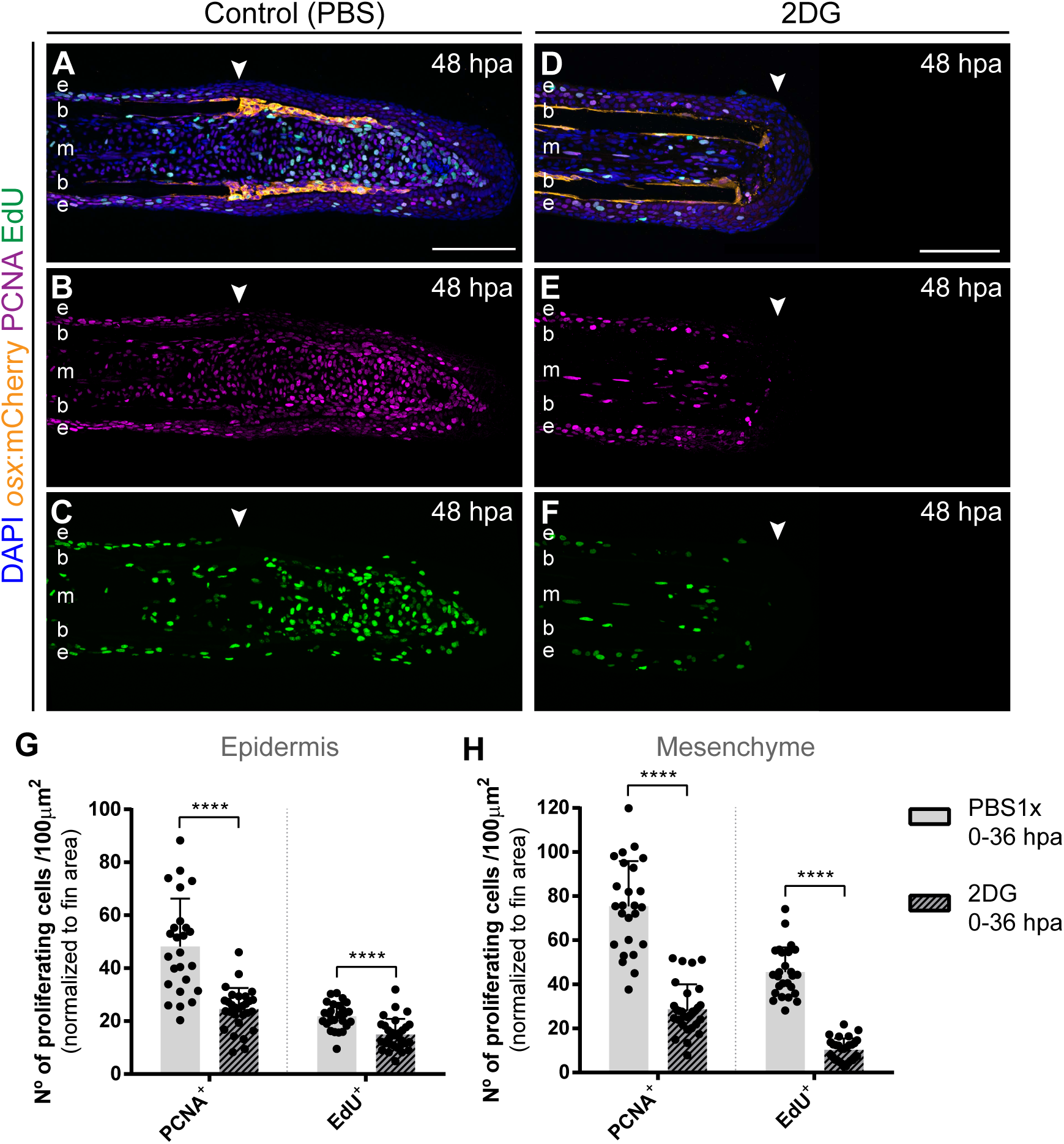
Inhibition of glycolysis impairs proliferation during blastema formation. (A-F) Representative 48 hpa cryosection images of osx:mCherry (orange) caudal fins immunostained for PCNA (magenta), EdU (green) and counterstained with DAPI (blue), in fish treated with (A-C) vehicle (PBS) or (D-F) 2DG. Arrowheads indicate amputation plane. Scale bar represents 100 µm. e: epidermis; b: bone; m: mesenchyme. (G-H) Quantification of the number of proliferative PCNA+ and EdU+ cells in the (G) epidermal and (H) mesenchymal compartments at 48 hpa, in caudal fins from controls and 2DG treated fish. Statistical analysis displayed on the graphs corresponds to Mann-Whitney test with Mean ± SD (n=25 (DMSO) and 29 (3PO) cryosections). ****P <0,0001.

**Supplementary Table 1:**
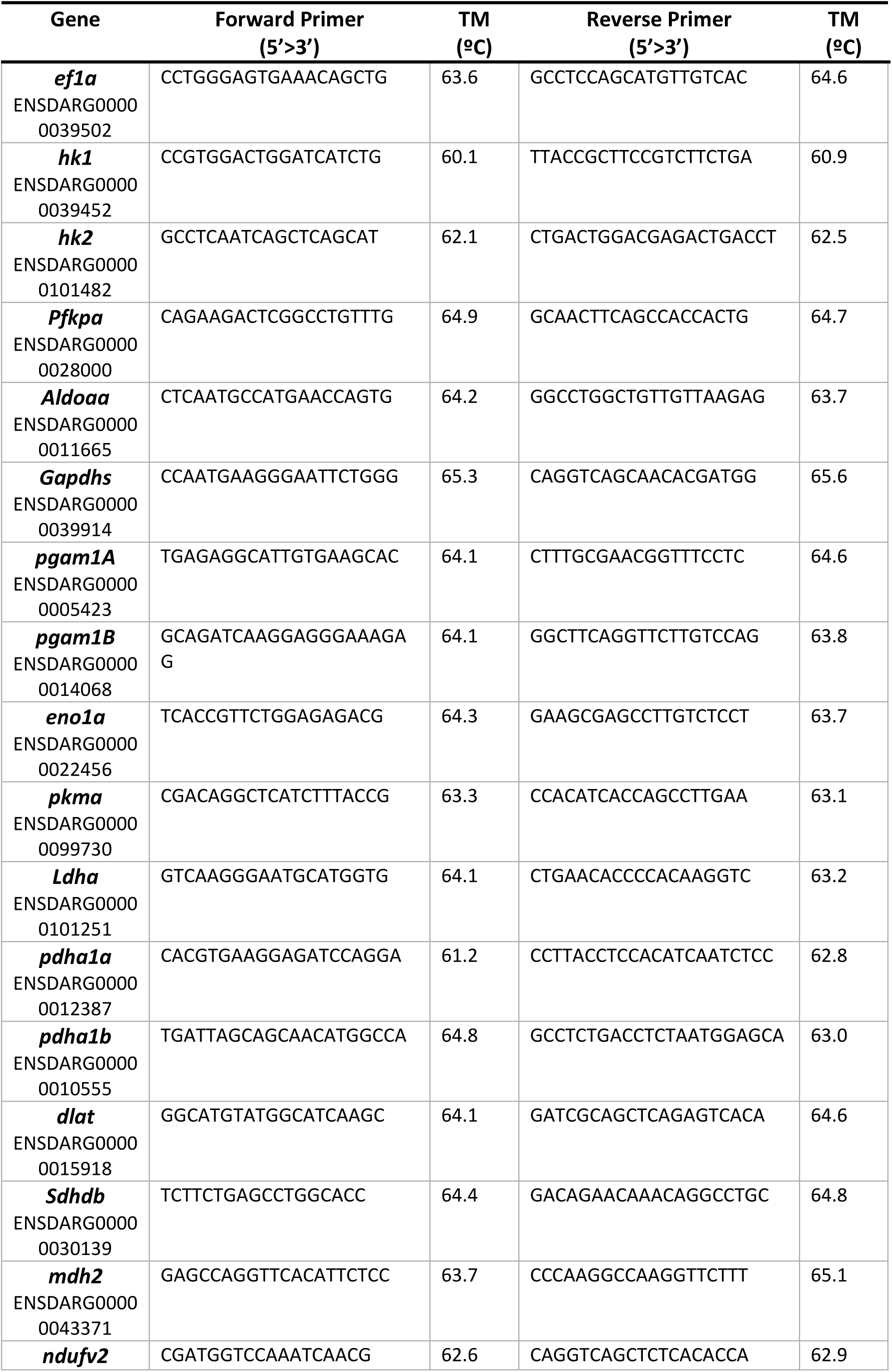
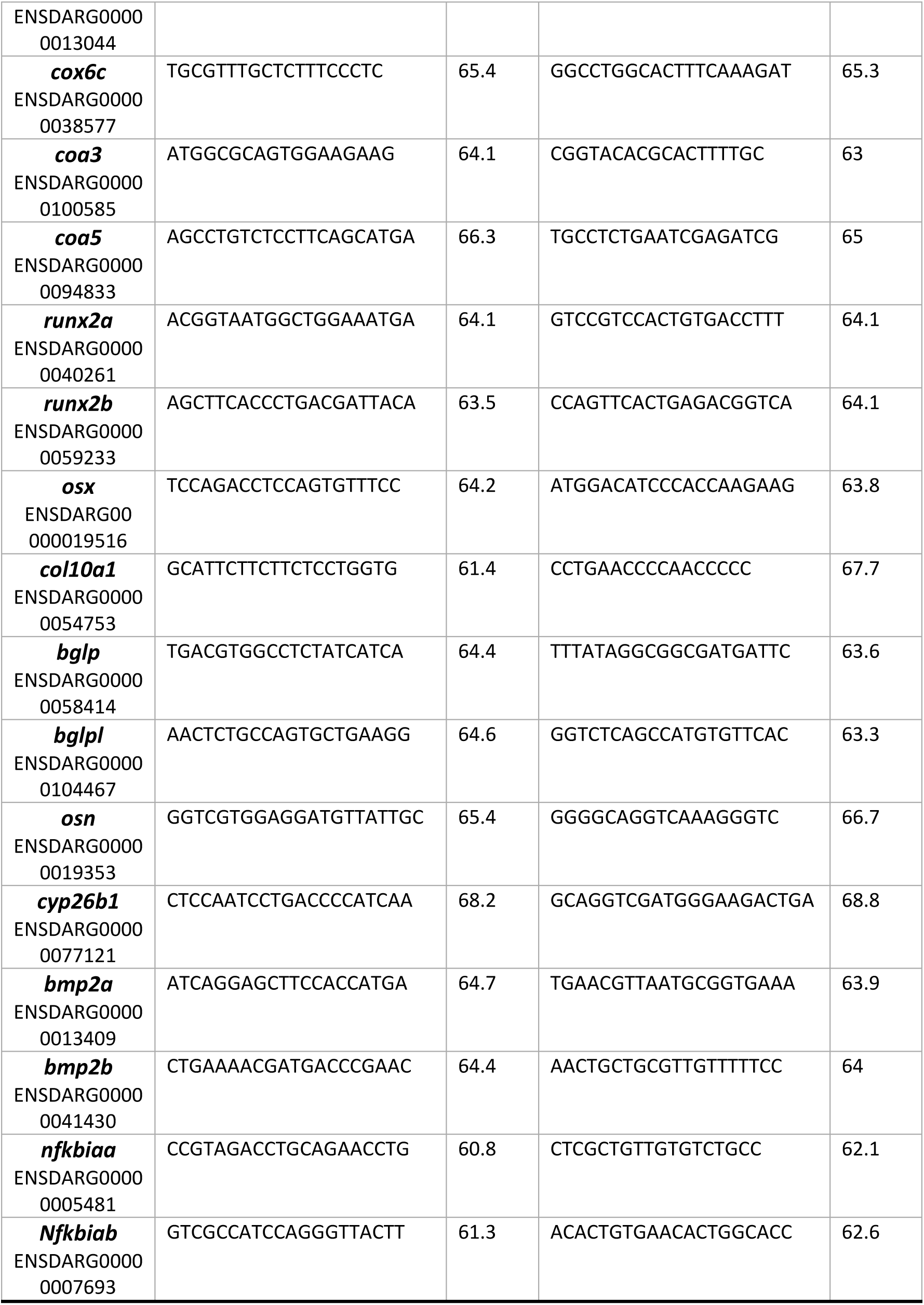
Primer sequences for q-PCR assay.

**Supplementary Table 2:**
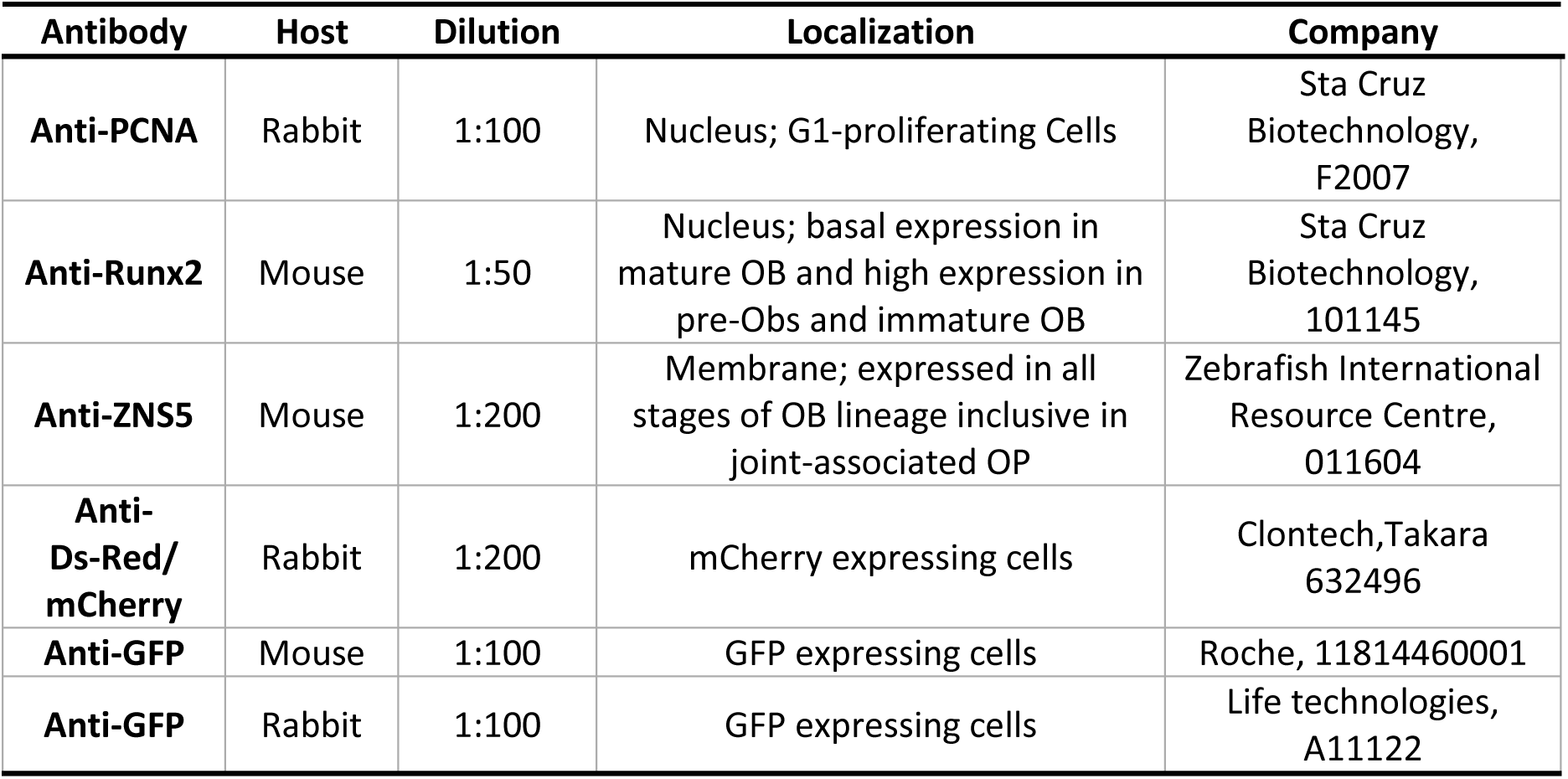
List of primary antibodies used for immunofluorescence assays.

**Supplementary Table 3:** List of primary antibodies used for immunofluorescence assays.

| Fluorophore | Host | Specificity | Dilution | Company |
| --- | --- | --- | --- | --- |
| Alexa Fluor 488 | Goat | Mouse | 1:500 | Invitrogen, A11001 |
| Alexa Fluor 488 | Goat | Rabbit | 1:500 | Invitrogen, A11070 |
| Alexa Fluor 568 | Goat | Mouse | 1:500 | Invitrogen, A11031 |
| Alexa Fluor 568 | Goat | Rabbit | 1:500 | Invitrogen, A11036 |
| Cy5 | Goat | Mouse | 1:250 | Invitrogen, A10524 |
| Alexa Fluor 647 | Donkey | rabbit | 1:500 | Jakson Imuno Research 711605152 |

**Supplementary Table 4:** List of sample size number and statistical test preformed for each quantitative experimental design.

| Figure | Sample size (n) | Statistical test |
| --- | --- | --- |
| Main Figures |  |  |
| **Fig 1B** | 4 cDNA pools per time-point | Paired t-test |
| **Fig 1D** | 0hpa= 18, 5hpa= 8, 10hpa= 16, 15hpa= 9, 20hpa= 9, 25hpa= 9 bony-rays from 3 different fish specimens per condition | Kruskal-Wallis for multiple comparisons |
| **Fig 1E** | 0hpa= 6, 6hpa= 10, 12hpa= 12, 18hpa= 9, 24hpa= 13 cryosections from 3 different fish specimens per time-point | Kruskal-Wallis for multiple comparisons |
| **Fig 1H** | 0hpa= 27, 12hpa= 26, 24hpa= 20 cryosections from 4-5 blastemas from 4 different fish specimens per time-point | Kruskal-Wallis for multiple comparisons |
| **Fig 1I** | 0hpa= 27, 12hpa= 26, 24hpa= 18 cryosections from 4-5 blastemas from 4 different fish specimens per time-point | Kruskal-Wallis for multiple comparisons |
| **Fig 2B** | 3 cDNA pools per time-point | t-test with Welch's correction |
| **Fig 2C** | 5 cDNA pools per time-point | Paired t-test |
| **Fig 2D** | 4 cDNA pools per time-point | Paired t-test |
| **Fig 2E** | 4 metabolite pools per time-point | Mann-Whitney |
| **Fig 2N** | 0hpa=59, 24hpa=83 bony-rays from 3 and 4 different fish specimens, respectively | Mann-Whitney |
| **Fig 3C** | 0 hpa: PBS=7, 2DG=8; 0-12hpa: PBS/2DG=7; 0-24hpa: PBS=4, 2DG=5; 0-36hpa: PBS=7, 2DG=5 fish per condition | Mann-Whitney |
| **Fig 3O** | 0-36hpa: PBS=13, SO=12 fish per condition | Mann-Whitney |
| **Fig 3S** | 0-36hpa: PBS:DMSO(1:1)=6, UK5099=5 fish per condition | Mann-Whitney |
| **Fig 4C** | 8 cDNA pools per condition | Paired t-test |
| **Fig 4F** | 0-12hpa: PBS=49, 2DG=69 cryosections from 5-6 blastemas from 6 different fish specimens per condition | Mann-Whitney |
| **Fig 4G** | 0-12hpa: PBS=49, 2DG=69 cryosections from 5-6 blastemas from 6 different fish specimens per condition | Mann-Whitney |
| **Fig 5E** | 0-12hpa: PBS=90, 2DG=82 bony-rays from 4 different fish specimens | Mann-Whitney |
| **Fig 5H** | 0-12hpa: PBS=23, 2DG=30 cryosections from 4-6 blastemas from 4 and 6 different fish specimens per condition, respectively | Mann-Whitney |
| **Fig 5I** | 0-12hpa: PBS=23, 2DG=27 cryosections from 4-6 blastemas from 4 and 6 different fish specimens per condition, respectively | Mann-Whitney |
| **Fig 5J** | 8 cDNA pools per condition | Paired t-test |
| **Fig 6E** | 0-36hpa: PBS=27, 2DG=25 cryosections from 4-6 blastemas from 6 and 5 different fish specimens per condition, respectively | Mann-Whitney |
| **Fig 6F** | 0-36hpa: PBS=27, 2DG=25 cryosections from 4-6 blastemas from 6 and 5 different fish specimens per condition, respectively | Mann-Whitney |
| **Fig 6G** | 0-36hpa: PBS=27, 2DG=25 cryosections from 4-6 blastemas from 6 and 5 different fish specimens per condition, respectively | Mann-Whitney |
| **Fig 6J** | 0-36hpa: PBS=28, 2DG=27 cryosections from 4-5 blastemas from 6 and 5 different fish specimens per condition, respectively | Mann-Whitney |
| **Fig 6K** | 0-36hpa: PBS=27, 2DG=27 cryosections from 4-5 blastemas from 6 and 5 different fish specimens per condition, respectively | Mann-Whitney |
| **Supplementary Figures** |  |  |
| **Sup Fig 2D** | 0-36hpa: DMSO=15, 3PO=14 fish per condition | Mann-Whitney |
| **Sup Fig 2H** | 0-36hpa: DMSO=4, MB6=4 fish per condition | Mann-Whitney |
| **Sup Fig 3G** | 0-12hpa: PBS=26, 2DG=35 cryosections from 4-6 blastemas from 5 and 6 different fish specimens per condition, respectively | Mann-Whitney |
| **Sup Fig 3H** | 0-12hpa: PBS=26, 2DG=35 cryosections from 4-6 blastemas from 5 and 6 different fish specimens per condition, respectively | Mann-Whitney |
| **Sup Fig 3O** | 24hpa: DMSO=10, 3PO=14 cryosections from 3-4 blastemas from 3 and 4 different fish specimens per condition, respectively | Mann-Whitney |
| **Sup Fig 4I** | 0-12hpa: PBS=17, 2DG=16 cryosections from 4-6 blastemas from 4 and 3 different fish specimens per condition, respectively | Mann-Whitney |
| **Sup Fig 4J** | 0-12hpa: PBS=17, 2DG=16 cryosections from 4-6 blastemas from 4 and 3 different fish specimens per condition, respectively | Mann-Whitney |
| **Sup Fig 4K** | 0-12hpa: PBS=21, 2DG=18 cryosections from 4-6 blastemas from 4 and 3 different fish specimens per condition, respectively | Mann-Whitney |
| **Sup Fig 4L** | 0-12hpa: PBS=21, 2DG=18 cryosections from 4-6 blastemas from 4 and 3 different fish specimens per condition, respectively | Mann-Whitney |
| **Sup Fig 5K** | 24hpa: DMSO=18, 3PO=19 cryosections from 3-5 blastemas from 4 and 5 different fish specimens per condition, respectively | Mann-Whitney |
| **Sup Fig 5L** | 24hpa: DMSO=18, 3PO=19 cryosections from 3-5 blastemas from 4 and 5 different fish specimens per condition, respectively | Mann-Whitney |
| **Sup Fig 6G** | 0-12hpa: PBS=25, 2DG=29 cryosections from 4-6 blastemas from 5 different fish specimens per condition | Mann-Whitney |
| **Sup Fig 6H** | 0-12hpa: PBS=25, 2DG=29 cryosections from 4-6 blastemas from 5 different fish specimens per condition | Mann-Whitney |

## Notes

### Competing Interest Statement

The authors have declared no competing interest.

